# Spatial Transcriptomics Unveils the Blueprint of Mammalian Lung Development

**DOI:** 10.1101/2025.04.05.647413

**Authors:** Mingyue Chen, Junjie Lv, Qiao Zhang, Qian Gong, Ting Zhao, Zhenping Chen, Haishen Xu, Nan Zhou, Shan Jiang, Jian Du, Xuepeng Chen, Yuwen Ke

## Abstract

Mammalian lung development is a complex, highly orchestrated process involving the precise coordination of diverse cell types. Despite significant advances, the spatial gene expression patterns and regulatory mechanisms within the developmental niches of different lung structures remain incompletely understood. In this study, we present a comprehensive spatial transcriptomic atlas of mouse lung development, spanning from the early pseudoglandular to the alveolar stage. We further uncover transcription factor (TF) regulation landscapes by integrating spatial epigenome, including novel TF-eRegulons critical for epithelial progenitors during early lung development. Our analysis also identifies hundreds of spatiotemporally dynamic cell-cell communications, such as *BMP8B*-mediated ligand-receptor signaling enriched in airway branching. Notably, we delineate the distinct developmental trajectories of alveolar AT1 and AT2 cells and reveal that collagen pathways facilitate their spatial convergence, forming primary alveoli during the canalicular-saccular transition. Together, this spatial transcriptomic atlas provides a foundational resource for understanding the complex cellular and molecular orchestration underlying mammalian lung development.

## Introduction

The lung is the primary organ responsible for gas exchange and tissue homeostasis in mammals [1]. Abnormal gene expression during lung development can lead to various respiratory diseases, including congenital pulmonary hypoplasia, pediatric asthma, and adult-acquired lung diseases [2–5]. Mammalian lung development involves a series of morphogenetic processes, such as airway branching, epithelial-mesenchymal interactions, vascularization, interstitial thinning, and alveolar formation [5–7].

Like in human, mouse lung developmental consists of five major stages based on morphological criteria: embryonic, pseudoglandular, canalicular, saccular, and alveolar stages [6]. During the embryonic stage, the two lung buds separate from the foregut, forming the primary lung airway. The airways then elongate and undergo branching morphogenesis during the pseudoglandular stage. The canalicular stage and saccular stages see the formation of the most distal airways, providing the foundational structure for future gas exchange. Notably, the saccular stage marks the end of branching morphogenesis and is characterized by ongoing alveolar epithelial differentiation, which is crucial for healthy pulmonary function after birth. Finally, alveolarization involves thinning of the alveolar septa and maturation of the microvascular network, culminating in the establishment of mature gas exchange.

Recent single-cell transcriptomic studies have identified various cell types in the developmental and adult lungs of mice and humans [8–14]. However, these studies often lack spatial information about the naïve locations of these cells. Investigating the spatial distribution of gene expression among different lung cell types, including epithelial, endothelial, and mesenchymal cells, is essential for understanding molecular microenvironments that drive the lung development.

Additionally, lung epithelial progenitor cells during the embryonic stage are typically distributed along the proximal-distal axis within the airways [15]. However, the chromatin accessibility landscapes of these critical progenitor cells remain largely unexplored. Furthermore, *in situ* epithelial-mesenchymal interactions within tissues play a vital role in branching morphogenesis and alveolar formation. Yet, the systematic identification of high-fidelity cell-cell communications and their regulatory roles in the formation of complex lung structures has not been fully resolved.

Spatially resolved transcriptomic methods provide both transcriptomic data and positional context for cells within tissues [16, 17]. Leveraging emerging high-resolution spatial transcriptomic technology DBiT-seq, we generated a comprehensive spatial transcriptomic atlas of mouse lung development to explore the dynamic spatial context of various cells and key molecules. We identified representative spatial lung cell clusters and validated spatial expression distributions of their marker genes. Additionally, we integrated spatial transcriptomic and epigenomic data to reveal relevant transcription factor (TF) regulation landscapes in lung epithelial progenitor cells. Notably, we also discovered novel spatiotemporal cell-cell communications that elucidate the spatial orchestration of lung development in mice. Overall, this study establishes a valuable spatial omics foundation for enhancing our understanding of mammalian lung development.

## Results

### Spatial transcriptomic atlas of lung development in mice

To systematically characterize spatial gene expression dynamics and cellular interactions during mammalian lung morphogenesis, we established a spatial transcriptomic atlas encompassing embryonic day 12.5 (E12.5) through postnatal day 14 (P14), capturing critical stages of lung development (Figures 1A, and S1A) [2]. Spatial transcriptomic (ST) profiling was performed using modified DBiT-seq [18] at 20 μm/10 μm resolution with biological replicates (Figures 1A, and S1A; Table S1). We used immunofluorescent (IF) staining to confirm mouse lung developmental stage progression, evidenced by NKX2.1+ epithelium localization and ACTA2+ smooth muscle differentiation patterns (Figure S1B). Following data preprocessing and quality control, our mouse lung ST atlas covered a total of 27,213 genes (Figure 1B; Table S2). The median unique molecular identifiers (UMIs) per ST spot was 5,365, with a range from 2,824 to 21,409 (Figure 1B). The median number of genes detected per spot reached 2,758 genes, with a range from 1,805 to 5,813 (Figure 1B).

**Figure 1.**
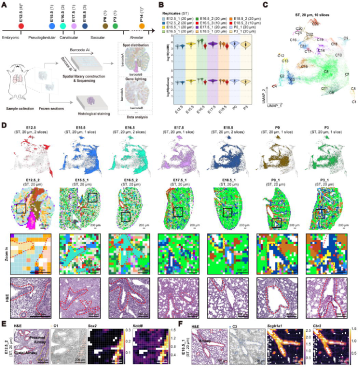
Spatiotemporal transcriptomic atlas of mouse lung development. **A**. Schematic workflow for constructing the spatial transcriptomic (ST) and epigenomic atlas during mouse lung development. Samples span five morphologically distinct developmental stages. The E12.5 data, marked with an asterisk (*), comprise two spatial transcriptomic libraries and two spatial ATAC-seq and RNA-seq libraries. The P14 ST data, also marked with an asterisk (*), were generated using Stereo-seq in this study. The number of biological replicates for each developmental time point is indicated in parentheses. **B**. Violin plots displaying the distribution of log10(nUMI) and log10(nGene) across samples. The resolution of the spatial transcriptomic (ST) data is indicated as 20 μm or 10 μm. **C**. UMAP embedding of all modified DBiT-seq profiled spots (20 μm resolution) across E12.5 to P3 stages. **D**. UMAP plots highlighting spots from different developmental time points (Row 1). Spatial visualization of one representative section for each time point (Row 2). Zoomed-in regions showing specific clusters (Row 3). Corresponding H&E staining of the adjacent slices for each section. Airway structures are delineated by red dashed lines (Row 4). Scale bars: 200 μm. **E**. Normalized spatial expression of C1 (Sox2+ epithelial progenitor cell-enriched) markers: *Sox2* (canonical) and *Kctd8* (newly identified) at E12.5. Red dotted lines indicate distal airways, and yellow dotted lines mark proximal airways. Scale bars: 200 μm. **F**. Normalized spatial expression of C3 (secretory cell-enriched) markers: *Scgb1a1* (canonical) and *Cbr2* (newly identified) at E18.5. Red dotted lines on H&E staining highlight airway structures. Scale bars: 200 μm.

Then, dimensionality reduction and unsupervised clustering of all 20 μm-resolution ST data (E12.5-P3) via uniform manifold approximation and projection (UMAP) revealed 24 transcriptionally distinct lung spatial clusters, each exhibiting unique molecular signatures and histo-topographical distributions (Figures 1C-D, and S1C-E; Table S3; Methods). Cluster-specific marker genes coincided very well with cluster spatial localization. For example, canonical epithelial progenitor marker *Sox2* and the novel candidate marker *Kctd8* showed co-localization in the early proximal airway (C1), while *Scgb1a1* (secretory cell marker) and *Cbr2* were specific to bronchial cluster C3 (Figures 1E-F). Additionally, *Wt1* expression characterized mesothelial cluster C14, and *Vegfc* marked vascular endothelial cluster C18 (Figures S1F-G).

Next, computational integration with existing single-cell RNA-seq references [10, 13] via cell2location [19] established high-confidence spatial annotations (Figures S2A-E). Spatial clusters showed predominant enrichment for important lung cell types, including Sox2+ epithelial progenitors (C1) and pulmonary neuroendocrine cells (PNECs; C4) (Figures S2C-E). Functional enrichment analysis of 21 out of 24 clusters (7 epithelial, 7 mesenchyme-derived, 7 vasculature-associated) uncovered cluster-specific biological programs, including ciliated cell specialization through cilium assembly machinery in C2 (Figures S2F).

For the subcellular resolution P14 Stereo-seq data in this study, we systematically evaluated spatial binning resolutions (10/20/25/50/100 μm) and identified bin100 (50 μm) as the optimal binning strategy, balancing gene detection sensitivity (median 1,984 genes/bin100) with spatial coherence (Figure S3A). Unsupervised clustering resolved eight spatial domains, with deconvolution confirming lung cell type associations (Figures S3B-C). The P14-C1 domain reflected lung bronchiolar architecture through spatial enrichment of club and ciliated cells (Figures S3C-E). In addition, limited immune cells were reported in reference atlases [13]. Our deconvolution analysis revealed associations between immune cell and spatial clusters in both DBiT-seq and Stereo-seq data (Figure S2D and S3C). However, the widely spread distributions of immune cells made it difficult to achieve clear spatial clustering (Figure S3E).

Overall, we generated a comprehensive spatial transcriptomic atlas of mouse lung development. Our findings demonstrate that this developing mouse lung ST atlas accurately reveals lung cell diversity and spatial gene expression patterns.

### Investigation of TF regulation networks in early developing mouse lungs by MISAR-seq

Spatial transcriptomic profiling at 20 μm resolution (E12.5_1 and E12.5_2) delineated proximal-distal segregation of epithelial progenitors (ST-C1: proximal-enriched; ST-C2: distal-enriched), which was also supported by single-cell transcriptomic deconvolution (Figures S4A-E). To elucidate the epigenetic regulatory landscape governing this spatial diversification of epithelial progenitors, we implemented a modified multiplexed *in situ* ATAC-RNA sequencing (MISAR-seq) platform [20] at 10 μm resolution for simultaneously mapping chromatin accessibility and gene expression in E12.5 lungs (SMO data) (Figures S5A-F, and S6A-B).

Chromatin accessibility profiling of E12.5 lungs demonstrated robust technical metrics, with 17,635 median high-quality fragments per spot for E12.5_3 and 19,339.5 fragments for E12.5_4 (Figures S5A-B; Table S2). Besides, we detected about a median 3,000 genes for spatial spots of E12.5 SMO data (Figure S5A; Table S2). Furthermore, unsupervised clustering of chromatin accessibility (ATAC-only, 9 clusters), transcriptome (RNA-only, 12 clusters), and integrated modalities (ATAC and RNA, 11 clusters) revealed a strong inter-modality concordance (Figures S5C-F; Methods).

Further integrative analysis identified 19,053 association links between chromatin accessibility peaks and target genes (P2G links), with associated heatmaps confirming the concordance for P2G links between chromatin accessibility and gene expressions (Figure 2A). We identified *Foxa2*, *Nkx2-1*, and *Foxf1* as principal transcriptional regulators through TF motif enrichment analysis, and subsequently mapped their binding sites across all P2G links (Figure 2A). While confirming the known *Etv5* and *Nkx2-1* regulatory interaction in epithelial populations [21], our analysis further identified novel associations involving *Ap1s3* with *Foxa2* motifs and *Lin7a* with *Nkx2-1* motifs (Figure 2A). In parallel, both *Myh11* and *Map3k20* showed enrichment in non-epithelial compartments for *Foxf1* TF motif (Figure 2A).

**Figure 2.**
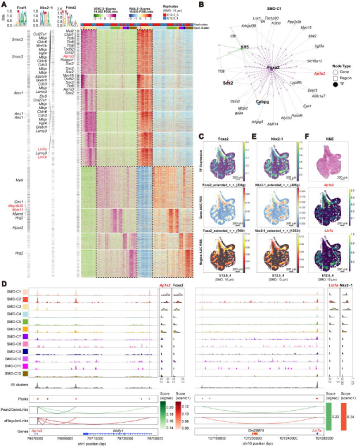
Spatial transcriptome and epigenome of the early developing mouse lung. **A**. Heatmap depicting chromatin accessibility and gene expression for 19,053 significant peak-gene linkages with E12.5_3 and E12.5_4 spatial multi-omic (SMO) data. Each row represents a peak-gene pair, with individual peaks potentially linked to multiple genes and vice versa. Peaks containing motifs for *Foxa2*, *Nkx2-1*, and *Foxf1* transcription factors (TFs) are annotated on the left. **B**. TF-region-gene regulatory network specific to SMO-C1. Dashed lines denote TF-region regulatory relationships, while solid lines represent region-gene regulatory interactions. **C**. Spatial mapping of *Foxa2* TF-eRegulon specific to spatial cluster SMO-C1 (Sox2+ epithelial progenitor-enriched), visualized through TF expression, Gene-AUC RSS, and Region-AUC RSS. Scale bars: 200 μm. **D**. Genome browser tracks illustrating chromatin accessibility (Top), peak loci, Signac-derived Peak2GeneLinks, SCENIC+ predicted eRegulonLinks (Middle), gene tracks (Bottom), and corresponding gene expression (right panel) for TF-eRegulon-target gene pairs: *Foxa2*-*Ap1s3* (Left) and *Nkx2-1*-*Lin7a* (Right). **E**. Spatial mapping of *Nkx2-1* TF-eRegulon specific to SMO-C3 (Sox9+ epithelial progenitor-enriched), displayed via TF expression, Gene-AUC RSS, and Region-AUC RSS. Scale bars: 200 μm. **F**. H&E staining of the adjacent slice of E12.5_4 and spatial visualization of target genes *Ap1s3* and *Lin7a* on E12.5_4, demonstrating their preferential localization along proximal and distal airway structures, respectively. Scale bars: 200 μm.

Furthermore, SCENIC+ TF regulatory module analysis which can integrate TF, enhancers and target genes [22] recapitulated known morphogenic circuits (*Foxa2*, *Gli1*, *Gata6-eRegulons*) [23–25] while uncovering 28 epithelial progenitor-specific TF-eRegulons (Figures 2B, and S6C-E). Spatial activation patterns mirrored important developmental knowledge: *Sox2-e*Regulon dominated proximal epithelial progenitors (SMO-C1), whereas *Etv5*/*Sox9*/*Nkx2-1* eRegulons characterized distal epithelial progenitors (SMO-C3) (Figures 2B, and S6B-C) [2]. Notably, we discovered many intriguing TF-eRegulons, such as the *Hnf1b, Grhl2*, and *Foxa2*, enriched in epithelial progenitors (Figures 2B, and S6B-E). *Hnf1b*, previously identified in human late tip epithelial cells [26], exhibited high gene expression in mouse SMO-C2 (stalk epithelial progenitors-enriched) (Figures S6B-D). We also found novel TF-eRegulons, such as *Rarb* in SMO-C1 and *Rbpjl* in SMO-C3 (Figures S6C, and S6E). *Rarb*, a retinoic acid (RA) nuclear receptor, has been implicated in regulating pulmonary functions [27], while *Rbpjl* is known to repress *Notch* target gene expression [28].

Extensive researches have established the roles of TFs in cell differentiation, niche formation, and regeneration during mouse lung development [2, 29]. Thus, we established a TF-enhancer-target gene regulatory paradigm to understand TF regulatory roles in the lung development. For example, *Foxa2*-eRegulon coordinated proximal/stalk progenitor maintenance (SMO-C1/C2), directly occupying *Ap1s3* enhancers to drive airway proximal-distal expression gradients (Figures 2C-D, 2F, S7A and S7E). Similarly, we demonstrated region-specific expression of *Nkx2-1* TF-eRegulon in distal progenitor (SMO-C3) (Figures 2D-E, and S7B). *Nkx2-1* is critical for normal morphogenesis and activates pulmonary differentiation-specific genes, including *Sftpa*, *Sftpb*, and *Sftpc* [30]. We also identified *Lin7a*, a target gene of *Nkx2-1*, concentrated in SMO-C3 filling the distal airway (Figures 2D, 2F, and S7E). *Lin7a* is recognized as a critical gene in the onset and progression of various diseases [31–34], suggesting a new role for *Lin7a* in early mouse lung development, potentially regulated by *Nkx2-1* binding to its enhancer (Figure 2D).

*Grhl2* is known to play a substantial role in lung branching morphogenesis, with alterations in *Grhl2* linked to bronchopulmonary dysplasia (BPD), a severe complication in premature infants [35, 36]. We observed markedly enhanced expression of *Grhl2* TF-eRegulon in proximal progenitors and stalk epithelial cells (SMO-C1/C2) (Figures S6C, and S7C). The expression of *Sox2*, *Foxa1*, and *Grhl2* was significantly enriched in the two clusters, consistent with previous findings (Figure S7D) [37, 38]. Furthermore, we identified *Sox2*, a well-known marker of lung proximal progenitor cells, as a target gene of *Grhl2* TF, highlighting the cooperative function of cell-specific TF-eRegulons in cell fate determination and maintenance (Figures S7D-E).

In addition to epithelial cells, we also explored TF regulons in mesenchymal and endothelial clusters (Figure S8A). As for non-epithelial cells, our analyses revealed canonical *Wt1* enriched in mesothelial SMO-C7 (Figure S8A). Additionally, we also identified novel regulators, like *Etv1* in proliferating matrix fibroblasts (SMO-C4) and *Zbtb20* in airway smooth muscle cell (ASMC; SMO-C6) (Figures S8A-B). The TF-eRugulon and peak patterns of *Foxf1* were significantly enriched around the proximal airway (Figure S8C). We further identified the *Foxf1*-*Map3k20* regulatory axis, with the *Foxf1* motif located in the enhancer region of *Map3k20* (Figure S8D). Notably, the spatial visualization of *Map3k20* mirrored the pattern of *Foxf1* TF (Figures S8C, and S8E). Finally, we demonstrated specific expression of *Map3k20* and *Foxf1* in mesenchymal SMO-C6 (Figure S8D), while the MAPK pathway in the lung mesenchymal layer has been shown to coordinate effective epithelial-mesenchymal interactions during lung development [39].

Our integrated spatial multi-omics data has systematically reconstructed the regulatory blueprint of pulmonary morphogenesis, revealing cell type-specific TF-eRegulon networks that consisted of master TF regulators, lineage-defining cis-regulatory elements, and functionally validated target genes in early mouse lungs.

### Spatial cell-cell communications in developing mouse lungs

During the lung development, cellular organization establishes distinct anatomical compartments including airway, vascular and mesenchymal niches. These spatially defined microenvironments and their associated signaling cascades critically regulate cellular fate determination and differentiation trajectories [40–42]. To systematically map intercellular communication networks, we employed SpatialDM, a computational framework specifically designed for spatial transcriptomic analysis incorporating spatial autocorrelation metrics [43], identifying hundreds of significant ligand-receptor (LR) interactions within developing lung niches (Figure S9A; Table S4). The calculated Moran’s I for these LRs demonstrated a strong positive correlation between biological replicates with Pearson’s R > 0.88 (Figure S9B). Introducing non-lung ST data from melanoma and intestine further revealed that LRs identified in the mouse lung displayed a clustered pattern distinct from melanoma and intestine ST data (Figure S9C).

Spatiotemporal analysis revealed distinct LR enrichment patterns across developing lung anatomical compartments, including mesenchyme, bronchi, alveoli, and vessels (Figure 3A). Previously reported LRs associated with WNT, NOTCH, and FGF signaling pathways [3, 44–46] exhibited compartment-specific enrichment. For instance, *FGF2*-*FGFR4*, *WNT2*-*FZD4*/*LRP5*, and *WNT5A*-*FZD4* were significantly enriched in the lung mesenchyme, while *BMP2*-*BMPR1B*/*ACVR2B*, *WNT3*-*FZD3*/*LRP6*, and *DLL3*-*NOTCH3* were more abundant in the bronchi (Figure 3A). *PTN*-*PTPRZ1* signaling was demonstrated with distinct enrichment in airway structures (Figure 3A), supporting PTN’s role in normal lung airway development [47]. We also observed *JAG1*-*NOTCH3* enrichment in vascular structures [48] (Figures 3A-B).

**Figure 3.**
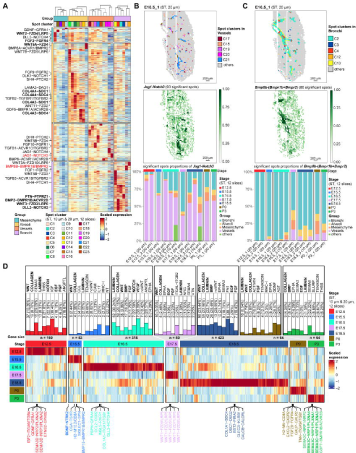
Spatial cell-cell communications in the developing mouse lung. **A**. Heatmap depicting lung structure-specific cell-cell communication dynamics during mouse lung development. All 20 μm-resolution ST spots were stratified into four anatomical groups: mesenchyme, alveoli, vasculature, and bronchi, based on structural localization. **B**. Spatial distribution of spot clusters within the vessel group (Top). Spatial local autocorrelation probability (1 - z_pval) map for *JAG1-NOTCH3* interactions (Middle). The histogram showing the proportion of *JAG1-NOTCH3* in different kinds of lung structure groups (Bottom). Scale bars: 200 μm. **C**. Spatial distribution of spot clusters within the bronchi group (Top). Spatial local autocorrelation probability (1 - z_pval) map for *BMP8B*-*BMPR1B*/*BMPR2* interactions (Middle). The histogram showing the proportion of *BMP8B*-*BMPR1B*/*BMPR2* interactions across lung structural groups (Bottom). Scale bars: 200 μm. **D**. Heatmap illustrating temporally dynamic ligand-receptor (LR) pairs during lung development. Overlaid bar plots (Top) indicate the pathway-specific distribution of identified LR interactions.

Furthermore, our analysis uncovered multiple novel LR interactions during murine lung development. Bone morphogenetic proteins (BMPs), like BMP2, BMP3, BMP4, BMP5, BMP6, and BMP7 are known to regulate mammalian lung development [49]. Notably, we identified BMP8B-mediated signaling as a novel regulator of airway morphogenesis, expanding its known roles in spermatogenesis and thermogenesis [50–52]. Spatial mapping revealed *BMP8B*-*BMPR1B*/*BMPR2* and *BMP8B*-*BMPR1B*/*ACVR2B* enrichment in the airway, particularly at branching points in the bronchi (Figures 3A, 3C, and S9D). The enrichment of *BMP8B*-*BMPR1B*/*ACVR2B* in bronchial structures kept a similar enrichment pattern during lung development, spanning from E12.5 to P14 (Figure S9D).

Besides, our temporal analysis for LR signaling also revealed stage-specific LRs. *BDNF*-*NTRK2* specifically enriched in the early stages of lung development, while *SEMA3C*-*NRP*/*PLXN* LR were enriched postnatally (P3) (Figure 3D). These LRs were enriched not only in canonical pathways (WNT, FGF, BMP, NOTCH) but also in COLLAGEN and LAMININ signaling (Figure 3D), implicating extracellular matrix remodeling in developmental regulation. Lung compartment enrichment change was observed for *COL1A2*-*ITGA11*/*ITGB1*, which transitioned from vascular-specific enrichment (E12.5) to dual airway-vascular distribution (E15.5–P3; Figure S9E), suggesting its dual role in vascular maturation and airway morphogenesis.

This comprehensive spatial LR analysis reveals hierarchical and dynamic LR signaling orchestrating pulmonary morphogenesis through known pathways and novel intercellular interactions. The spatiotemporal enrichment redistributions of LRs demonstrate the plasticity of cell-cell communications during lung development, offering new insights into developmental signaling.

### Temporal development of lung conducting airway and vascular cells

To delineate the spatial organization of pulmonary conducting airways, we performed sub-clustering analysis on 2,233 spatially resolved airway spots from developing mouse lungs (Figures S10A-B; Methods). This analysis identified five distinct transcriptional clusters: AW-C1 (Sox2+ early epithelium-enriched), AW-C2/C3 (secretory/ciliated cell-enriched), AW-C4 (neuroendocrine cell-enriched), and AW-C5 (ASMC-enriched) (Figure S10C). Spatial mapping confirmed compartment-specific expression of canonical markers: *Spag17* (ciliated cells), *Scgb3a2* (secretory cells), *Pcsk1* (neuroendocrine cells), and *Myh11* (ASMCs) within conducting airway structures (Figure S10D). Temporal comparison revealed dynamic proportion shifts among these epithelial subpopulations throughout pulmonary morphogenesis (Figure S10E). PNECs, representing a rare yet functionally critical epithelial population, play essential roles in pulmonary physiology through bioactive amine and neuropeptide production, with established links to asthma pathogenesis and small cell lung cancer development [53–55]. Our pseudotime analysis showed that PNECs emerged very early in the conducting airway (Figures S10F-G), which was consistent with previous reports [56, 57].

Vascular compartment analysis identified six vessel-associated subclusters, such as arterial endothelial cells (AECs)-enriched EC-C2 and pericyte-enriched EC-C5 populations (Figures S11A-B). Molecular characterization showed *Trpc6* as a definitive pericyte marker (EC-C5) and *Kit* as a general capillary (gCap) signature (EC-C6) (Figure S11C). Near-cellular resolution spatial mapping (10 μm) revealed the scattered expression of *Kit* and *Trpc6* in regions outside of large airways and vascular structures at 10 μm-resolution E18.5 ST sections (Figure S11D).

In summary, our spatial transcriptomic analysis delineates the molecular characteristics of conducting airway and vascular cells in mouse lungs, confirming PNECs as an early specified epithelial lineage during pulmonary morphogenesis.

### Divergent developmental trajectories of alveolar epithelial lineages

The alveolar compartment, comprising specialized epithelial cells, fibroblasts, and capillary networks, constitutes the fundamental gas exchange unit in pulmonary architecture [5]. To elucidate alveolar development, we integrated spatial transcriptomic datasets of alveolar cells from E16.5 and E18.5 lungs with Sox9+ distal progenitors from E12.5 lungs, identifying eight distinct clusters encompassing alveolar epithelial populations and progenitor states (Figures S12A-B). Spatiotemporal analysis revealed progressive maturation signatures, with AT1 marker *Ager* and AT2 maker *S*ftpa1 exhibiting significant gene expression increases during the canalicular-to-saccular transition (Figure 4A).

**Figure 4.**
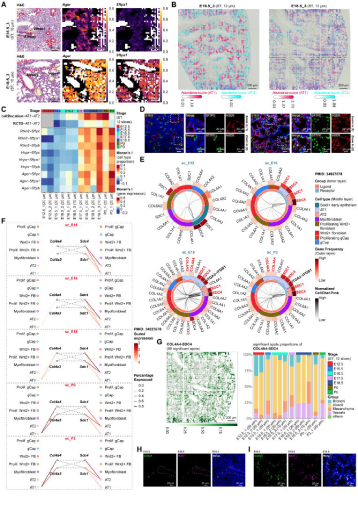
Spatial convergence of AT1 and AT2 cells and alveolar niche formation. **A**. High-resolution (10 μm) spatial transcriptomic profiles of AT1 marker *Ager* and AT2 marker *S*ftpa1 expression at E16.5 and E18.5. Red dashed lines demarcate airway structures, while yellow dashed lines indicate vascular networks. Scale bars: 200 μm. **B**. Spatial distribution patterns of AT1 and AT2 cells visualized through cell2location abundance scores at E16.5 and E18.5. Scale bars: 200 μm. **C**. Heatmap comparing spatial autocorrelation (Moran’s I) between RCTD weights and cell2location scores for AT1 and AT2 cells, alongside established AT1 (*Rtkn2*, *Hopx*, *Ager*) and AT2 (*Sftpc*, *S*ftpa1, *Sftpb*) marker gene pairs. **D**. Immunofluorescence staining of AGER (AT1) and SFTPC (AT2) at E16.5 and E18.5 lungs. scale bars: 50 μm. The zoomed-in regions are showed on the right. scale bars: 20 μm. **E**. Circle plot illustrating LRs involving distal epithelial cell populations, including Sox9+ early epithelium, AT1, and AT2 cells, as receptor-expressing populations in collagen signaling pathways, derived from scRNA-seq data (PMID: 34927678). **F**. Expression probability and level for collagen ligands (*COL4A4*/*COL4A3*) and their receptors (*SDC4*/*SDC1*) across alveolar niche cell subtypes in the scRNA-seq data (PMID: 34927678). Prolif. gCap: proliferating gCap; Wnt2+ FB: Wnt2+ fibroblast; Prolif. Wnt2+ FB: proliferating Wnt2+ fibroblast. **G**. Spatial probability map (1 - z_pval) of *COL4A4-SDC4* co-localization (Left) with corresponding structural distribution analysis across lung compartments (Right). **H-I**. RNA fluorescence in situ hybridization (FISH) confirming the spatial expression of *Col4a4* and *Sdc4* at E16.5 **(H)** and E18.5 **(I)**, with white dashed lines outlining developing airway structures. Scale bars: 20 μm.

Furthermore, STREAM pseudotime analysis [58] delineated divergent differentiation paths for alveolar epithelial cells: Sox9+ progenitors (AL-C1) segregated into AT1 precursors (AL-C2) and AT2 precursors (AL-C3) through distinct developmental paths (Figure S12C), supporting the paradigm of AT1/AT2 lineage-independent specification rather than a common progenitor model [8, 59–61]. RNA velocity analysis using scVelo [62] independently validated these segregated differentiation paths, while maintaining the plasticity of AT2 cells (AL-C5) to transdifferentiate into AT1 lineages (AL-C4) (Figures S12C-D). It was consistent with previous reports [63, 64]. AT1 and AT2 lineage-specific gene signatures were also identified: *Sftpb*, *Sftpc*, *Meg3*, *Lamp3*, and *Etv5* for the AT2 lineage, and *Rtkn2*, *Hopx*, *Ndnf*, *Sema3a*, and *Ager* for the AT1 lineage (Figures S12E-F).

Altogether, our spatial transcriptomic data supported distinct developmental paths for AT1 and AT2 cells during the canalicular-to-saccular transition in mouse lungs.

### The spatial gathering of AT1 and AT2 cells and alveolar niche formation

In addition to their distinct developmental trajectories, AT1 and AT2 cells exhibited dynamic spatial location reorganization during alveolar morphogenesis. According to 10 μm-resolution spatial transcriptomic data in E16.5 and E18.5 lungs, our results revealed AT1/AT2 segregated spatial distributions in E16.5 canalicular stage lungs would transition to a co-localized pattern at E18.5 saccular stage lung (Figure 4B). Spatial autocorrelation analysis using RCTD and cell2location algorithms demonstrated a progressive AT1-AT2 proximity, evolving from dispersed distributions before E16.5 to coordinated localization from E17.5 onwards (Figure 4C). The correlation analysis of cellular abundance scores confirmed this spatial location reorganization, with a negative correlation at E16.5 shifting to positive correlation at E18.5 (Figure S13A). Immunofluorescence (IF) staining and RNA FISH using AT1 marker *Ager* and AT2 marker *Sftpc* both confirmed the ST observation of AT2 cell dispersion along airway tips at E16.5 and subsequent alveolar co-localization with AT1 cells by E18.5 (Figures 4D, and S13B).

The spatial convergence of AT1 and AT2 cells signifies alveolar niche establishment, mediated by specialized cell-cell communication networks. Previous studies implicated collagen/integrin-mediated adhesion and actin cytoskeleton remodeling in alveolar elongation and mesenchymal migration [65–67]. We integrated ST and scRNA-seq data [10] and identified a notably dynamic collagen/integrin signaling pathways orchestrating niche formation and cell communications within alveolar compartments (Figures 3A, 4E, and S13C-E), such as *COL1A1*/*2*-*ITGA3*/*ITGB1* LRs in myofibroblast-AT1 communications and *COL4A1*/*2*-*ITGA3*/*ITGB1* LRs in capillary-AT1 communications (Figures 4E, and S13C). Additionally, we found *ITGA2/ITGB1*-mediated LR signaling such as *COL1A1*/*2*-*ITGA2*/*ITGB1*, *COL4A1*/*2*-*ITGA2*/*ITGB1*, and *COL6A1*/*2-ITGA2*/*ITGB1* LRs enriched in mesenchyme-capillary communications (Figures S13C, and S13E).

Notably, *COL4A3*/*4*-*SDC1* and *COL4A3*/*4*-*SDC4* LR interactions, which are part of the *SDC* receptor-mediated collagen pathways crucial for cell adhesion [68], exhibited the enriched specificity between alveolar epithelial cells, but were absent in alveolar endothelial and mesenchymal cell communications (Figures 4E, and S13C). scRNA-seq data [10] revealed the temporal upregulation of *COL4A3/4* in AT1 cells (E16-E18), and relatively stable *SDC1/4* expression in AT2 cells (Figure 4F). ST visualization corroborated these findings (Figure 4G). We then applied RNA FISH and IF staining, and further validated indeed *Col4a4* and *Sdc4* enrichment in distal airway at E16.5, and alveolar structures at E18.5 (Figures 4H-I, and S13G-H). We thus proposed that these type IV collagen-mediated signaling were relevant for the spatial gathering of AT1 and AT2 cells.

This integrated analysis revealed that alveolar AT1 and AT2 cells underwent spatial gathering during the canalicular-saccular transition, which was probably mediated by type IV collagen*-SDC* receptor signaling.

### Dynamic gene expression of alveolar domains during perinatal transition in mouse

The transition from intrauterine hypoxia to postnatal oxygenation represents a critical phase in mammalian lung development, requiring precise molecular reprogramming to establish gas exchange competence [1, 69]. To resolve these underlying molecular adaptations, we performed dynamic gene expression analysis for alveolar ST domains during perinatal transition including prenatal stage (E18.5), birth (P0), and early postnatal (P3) stages (Figure S14A, Methods).

Differential expression analysis revealed progressive downregulation of hypoxia-inducible factor *Hif3a* (a hypoxia-inducible factor), which was consistent with diminished hypoxic signaling as autonomous pulmonary oxygenation postnatally (Figure S14B). This temporal gene dynamic is particularly important, given the established role of *Hif3a* in alveolarization and distal epithelial differentiation for maintaining normal alveolar-vascular architecture [70, 71]. The down-regulated genes further suggested their involvement in *Wnt*-mediated cell signaling and cellular hypoxia response during environmental oxygen adaptation (Figure S14C).

Additionally, *Tgfbi*, a known regulator of NF-κB-dependent angiogenesis during early alveolarization [72], exhibited stage-specific induction, transitioning from near-undetectable levels at E18.5 to markedly up-regulated levels by P3 in alveolar domains (Figure S14B). Functional enrichment analysis connected these temporally up-regulated genes to critical biological processes, including epithelial-mesenchymal crosstalk and NF-κB transcriptional activation, highlighting their coordinated roles in alveolar maturation and microvascular development (Figure S14C). ST visualization further confirmed *Hif3a*, *Fgf10*, *Fn1*, and *Rtkn2* were highly enriched in the prenatal hypoxic alveolar zone, while *Tgfbi* and *Bmpr2* dominated in alveolar regions after the birth (Figures S14D-E).

Overall, our results above delineates the transcriptomic transition during perinatal adaptation in mouse lungs.

### Spatiotemporal expressions of transposable elements during mouse lung development

Protein-coding regions constitute merely ∼2% of the mammalian genome. Millions of transposable elements (TEs) are interspersed throughout the whole genome, but their functional roles remaining enigmatic [73–75]. Although TE activity is predominantly silenced in somatic tissues, yet certain elements maintain transcribed in specific biological contexts, such as neural development and early embryogenesis. The spatial expression dynamics of TEs during lung development, however, remain unexplored.

Next, our spatial transcriptomic analysis revealed active TE expression across developing lung compartments (Figure 5A). Notably, L1 elements like L1Md-F2, L1-Mus1, and L1Med elements exhibited pronounced transcription activity in progenitor populations, including epithelial and endothelial progenitors (Figure 5A). We also observed cluster-specific TE transcription activity patterns. For example, RLTR4-MM-int demonstrated airway epithelial specificity (C2/C3), while RLTR6-Mm showed mesothelial enrichment (C14) (Figures 5B-C).

**Figure 5.**
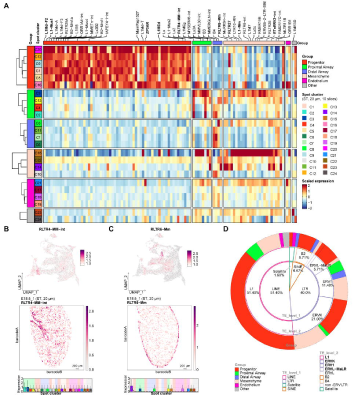
Spatiotemporal expression of transposable elements during the mouse lung development. **A**. Heatmap showing the scaled expression level of transposable elements (TEs) during the mouse lung development in different lung groups. All the 20 μm-resolution ST spots are divided into six groups based on lung structures and cell states: progenitor, proximal airway, distal airway, mesenchyme, endothelium, and others. **B-C**. UMAP visualization of normalized TE expression levels (Top). Spatial highlighting of normalized TE expression levels (Middle). Box plots showing the normalized TE expression levels across different clusters (Bottom). **(B)** RLTR4-MM-int and **(C)** RLTR6-MM. Scale bars: 200 μm. **D**. Multi-tiered pie chart delineating the compositional distribution of expressed TEs in developing murine lungs.

Classification of differentially expressed TEs identified long interspersed nuclear elements (LINEs) and long terminal repeats (LTRs) as the predominant classes (>90% combined) in the developing mouse lung (Figure 5D). By contrast, short interspersed nuclear elements (SINEs) representing ∼7% of active elements (Figure 5D). Specifically, the endothelial clusters exhibited enrichment only for Satellite and L1 elements, whereas both proximal and distal epithelium showed co-expression of ERV1 retrotransposons. (Figure 5D).

These TE spatial expression analyses show that extensive active TE expression exists throughout mouse lung development, with L1 and ERV elements emerging as the predominant classes exhibiting dynamic expressions across different pulmonary structures.

## Discussion

The mammalian lung is a major respiratory organ whose development involves the progressive specialization of diverse cellular lineages through precisely orchestrated morphogenetic processes. This study establishes a comprehensive spatial transcriptomic atlas encompassing five critical stages of pulmonary development, providing novel and detailed insights into the spatiotemporal signaling networks governing pulmonary linage maturation. Building upon recent studies [39, 63, 76, 77], we systematically mapped LR interactions across bronchial, alveolar, vascular, and mesenchymal compartments and delineated cellular responses to developmental signals during lung organogenesis. Beyond canonical WNT, FGF, and BMP signaling pathways, we identified novel LR interactions, including *BMP8B*-*BMPR1B*/*BMPR2* in proximal airways and *JAG1*-*NOTCH3* in vascular structures, expanding our understanding of pulmonary developmental signaling networks.

Additionally, spatial pseudotime analysis revealed that Sox9+ progenitors (AL-C1) diverge into distinct AT1 (AL-C2) and AT2 (AL-C3) precursor populations before maturing into specialized AT1 (AL-C4) and AT2 (AL-C5) lineages. Notably, while AT2 cells initially localize to airway tips at E16.5, they subsequently converge with AT1 cells in alveolar niches by E18.5, a process potentially mediated by *COL4A3*/*4*-*SDC1* and *COL4A3*/*4*-*SDC4* interactions, as supported by scRNA-seq, ST data, IF staining and RNA FISH. Our findings extended previous work demonstrating AT1 cells as signaling hubs during perinatal maturation [13, 67].

While this high-resolution spatial transcriptomic and epigenomic atlas provides significant insights, several technical limitations warrant consideration. Current 10 μm-resolution spatial-omic platforms cannot achieve real single-cell resolution, resulting in the incomplete detection of immune cells and alveolar capillary (aCap) populations that are readily identifiable in parallel single-cell datasets. To address this limitation, we implemented rigorous spatial-single cell integration strategies to improve ST cluster annotations and biological interpretations, complemented by immunofluorescence and RNA FISH validation. Additionally, the complexity of lung architecture and the limited availability of mesenchyme-specific markers constrained deeper analysis of stromal populations, necessitating higher-resolution spatial technologies and lineage-tracing approaches for comprehensive characterization.

## Conclusion

This study establishes a comprehensive spatial transcriptomic atlas spanning critical phases of murine pulmonary development, from late embryonic stages to alveolar stage. We delineated the TF regulatory networks governing early lung morphogenesis by identifying both novel and known TF-eRegulons. Our analysis identified hundreds of spatiotemporally dynamic cell-cell communications via LR signaling, providing mechanistic insights into lung compartment-specific regulatory programs. Notably, we demonstrated that AT1 and AT2 cells, which originated from separate precursors, underwent spatial convergence through type IV collagen-SDC mediated signaling pathways during the canalicular-saccular transition. Additionally, our investigation of transposable element dynamics revealed persistent expression of L1, ERVK, and ERVL-MalR elements in progenitor and mesenchymal populations. Collectively, this study delineates the spatial gene expression landscapes and regulatory networks governing lung compartment-specific development, providing an essential spatial multi-omics resource for understanding of mammalian pulmonary development.

**Table 1.** PCR Oligo and DNA Barcode Sequences for spatial RNA-seq and ATAC-seq library preparation.

**Table 2.** The data summary of spatial transcriptome in developing mouse lungs.

**Table 3.** Identified marker genes for ST spot clusters in mouse lungs.

**Table 4.** Significantly enriched ligand-receptor pairs according to spatial transcriptome of developing mouse lungs.

**Table 5.** The sequences of RNA FISH probes.

## Materials and methods

This is available in File S1.

## Ethical statement

Procedures were approved by the Institutional Animal Care and Use Committees of Guangzhou National Laboratory (Animal Ethics Approval Number GZLAB-AUCP-2022-10-A05).

### Data availability

All spatial data from this study are available in GSA database with the accession number CRA014010 (https://ngdc.cncb.ac.cn/gsa/s/bJ3k9E1z).

### Code availability

All code used for data analysis at GitHub (https://github.com/EddieLv/Temporal-and-spatial-development-of-mouse-lung), and BioCode at the National Genomics Data Center, Beijing Institute of Genomics, Chinese Academy of Sciences / China National Center for Bioinformation (BioCode: BT007890, https://ngdc.cncb.ac.cn/biocode/tool/BT007890).

### CRediT author statement

**Mingyue Chen**: Data curation, Formal analysis, Investigation, Methodology, Validation, Visualization, Writing – original draft. **Junjie Lv**: Data curation, Formal analysis, Investigation, Methodology, Software, Visualization, Writing – original draft. **Qiao Zhang**: Data curation, Methodology. **Qian Gong**: Investigation. **Ting Zhao**: Methodology. **Zhenping Chen**: Investigation. **Haishen Xu**: Data curation. **Nan Zhou**: Investigation. **Shan Jiang**: Methodology. **Jian Du**: Conceptualization. **Xuepeng Chen**: Conceptualization, Formal analysis, Funding acquisition, Methodology, Project administration, Resources, Supervision, Writing – review & editing. **Yuwen Ke**: Conceptualization, Formal analysis, Project administration, Supervision, Writing – review & editing.

## Acknowledgments

This work was supported by grants from the startup funding of Guangzhou Laboratory, the Major Project of Guangzhou National Laboratory (No. GZNL2023A03005) and National Natural Science Foundation of China (32170600).

## Supplementary material

**Figure S1.**
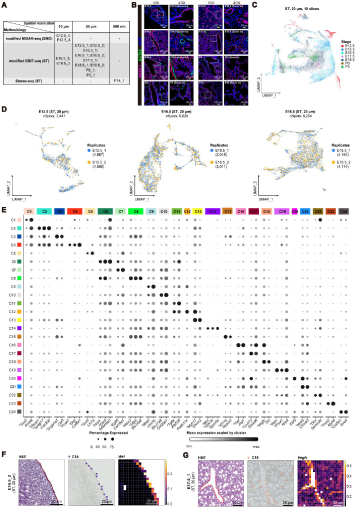
Data quality and characterization of the spatial transcriptomic atlas during mouse lung development. **A**. Summary of the resolution and spatial library construction methods for each tissue section. **B**. Representative immunofluorescence staining images for the epithelial cell marker NKX2.1 and the smooth muscle cell marker ACTA2 in developing mouse lungs from E12.5 to P14. Scale bars: 100 μm. The zoomed-in regions are showed on the right. scale bars: 20 μm. **C**. UMAP visualization of all modified DBiT-seq profiled spots (20 μm resolution) stratified by developmental stages. **D**. UMAP visualization of E12.5 (Left), E16.5 (Middle), and E18.5 (Right) modified DBiT-seq profiled spots (20 μm resolution) stratified by biological replicates. **E**. Dot plot illustrating the normalized gene expression levels across 24 spatial clusters. Dot size represents the percentage of expressed genes. Genes marked with an asterisk (*) are canonical markers, while the remaining genes are newly identified in this study. **F-G**. Normalized spatial gene expression patterns of cluster marker genes: *Wt1* **(F)** and *Vegfc* **(G)** on ST sections. Scale bars: 200 μm.

**Figure S2.**
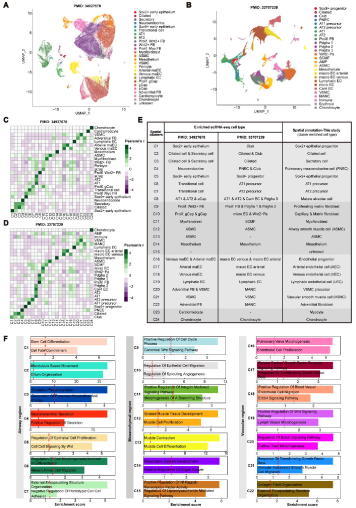
Single-cell mapping and functional enrichment analysis of spatial clusters. **A-B**. UMAP embeddings of the two published single-cell RNA-seq datasets (PMID: 34927678 and 33707239). **C-D**. Heatmaps displaying the distribution of scaled cell type abundances across spatial clusters for the two single-cell RNA-seq datasets (PMID: 34927678 and 33707239). **E**. Table summarizing enriched single-cell RNA-seq cell types for the 24 spatial clusters (20 μm-resolution data) and the manually curated spatial annotations used in this study. **F**. Bar plots of Gene Ontology (GO) functional enrichment for sub-clusters of airway, mesenchymal, and vascular regions. The red dotted line indicates the significance cutoff (p-value = 0.05).

**Figure S3.**
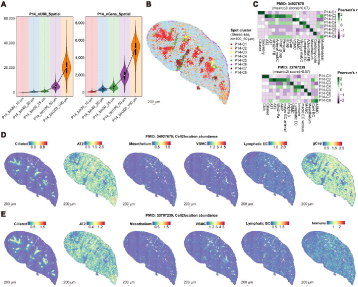
Spatial annotation and deconvolution analysis of the subcellular-resolution Stereo-seq data. **A**. Violin plots displaying the distribution of nUMI and nGene across different resolutions of P14 Stereo-seq data. Bin100 (50 μm) was selected for downstream analyses. **B**. Spatial visualization of P14 data, annotated based on unsupervised clustering results. Scale bars: 200 μm. **C**. Heatmaps illustrating the distribution of scaled cell type abundances across spatial clusters for two single-cell RNA-seq datasets (PMID: 34927678 and 33707239). **D**. Spatial mapping of single-cell RNA-seq-derived cell type abundances (ciliated, AT2, mesothelium, VSMC, lymphatic EC, and gCap; PMID: 34927678) on P14 data using cell2location. Scale bars: 200 μm. **E**. Spatial mapping of single-cell RNA-seq-derived cell type abundances (ciliated, AT2, mesothelium, VSMC, lymphatic EC, and immune cells; PMID: 33707239) on P14 data using cell2location. Scale bars: 200 μm.

**Figure S4.**
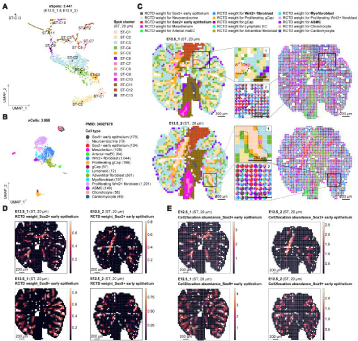
Spatial transcriptomic data visualization and spot-cluster deconvolution during early murine lung development. **A**. Re-clustering UMAP of 20 μm-resolution E12.5 ST data (E12.5_1 and E12.5_2), revealing 13 distinct spot clusters. **B**. UMAP visualization of reanalyzed E12 single-cell RNA-seq data (PMID: 34927678). **C**. Spatial visualization of re-annotated spot clusters at E12.5 (Left). Deconvolution using RCTD to map cell type proportions from single-cell RNA-seq data onto each spot (Right). Zoomed-in views of airway regions for the E12.5_1 and E12.5_2 ST sections (Middle). Scale bars: 200 μm. **D**. Spatial mapping of deconvoluted RCTD weights for Sox2+ early epithelium and Sox9+ early epithelium on the two replicate E12.5 ST sections. Scale bars: 200 μm. **E**. Spatial mapping of cell2location abundance scores for Sox2+ early epithelium and Sox9+ early epithelium on the two replicate E12.5 ST sections. Scale bars: 200 μm.

**Figure S5.**
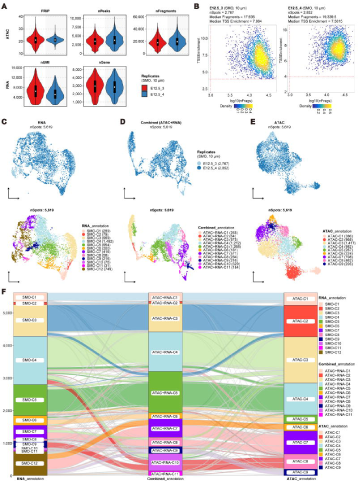
Quality control, clustering, and mapping results of ATAC-seq and RNA-seq data from E12.5 spatial multi-omics. **A**. Violin plots displaying FRiP, nPeaks, nFrags, nUMI, and nGene across two 10 μm-resolution E12.5 slices (E12.5_3 and E12.5_4). **B**. Density plots illustrating the distribution of log10(nFrags) and TSS enrichment scores for E12.5_3 (Left) and E12.5_4 (Right). **C-E**. Re-clustering UMAP of two E12.5 slices, with spots labeled by replicates (Top) and cluster annotations (Bottom), separately for RNA-seq **(C)**, combined data **(D)**, and ATAC-seq **(E)**. **F**. Sankey plot demonstrating the correlation between RNA-seq, ATAC-seq, and combined data annotations.

**Figure S6.**
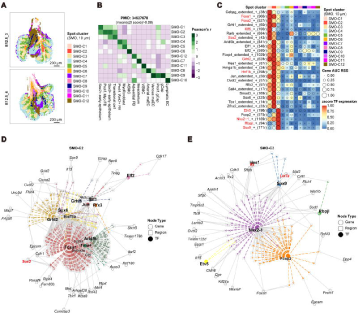
Gene regulatory networks of epithelial clusters in early mouse lung development. **A**. Spatial visualization of RNA clusters on E12.5_3 and E12.5_4 sections using modified MISAR-seq data. Scale bars: 200 μm. **B**. Heatmaps displaying the distribution of scaled cell type abundances across spatial clusters for the single-cell RNA-seq dataset (PMID: 34927678). **C**. Heatmap/dot plot illustrating the z-score TF expression (color scale) and cell-type specificity (RSS, size scale) of TF-eRegulons enriched in epithelial clusters. **D-E**. TF-region-gene regulatory networks for SMO-C2 **(D)**, and SMO-C3 **(E)**. Dashed lines represent TF-region regulatory relationships, while solid lines indicate region-gene regulatory interactions.

**Figure S7.**
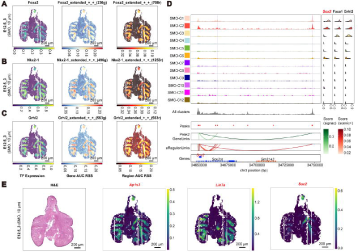
Spatial visualization of epithelial clusters-enriched transcription factors and their target genes during early mouse lung development. **A-C**. Spatial mapping of spot cluster-specific TF-eRegulons, showing TF expression, Gene AUC RSS, and Region AUC RSS for **(A)** SMO-C1 (Sox2+ epithelial progenitors-enriched; *Foxa2*), **(B)** SMO-C3 (Sox9+ epithelial progenitors-enriched; *Nkx2-1*), and **(C)** SMO-C1/C3 (Sox2+ and stalk epithelial cells-enriched; *Grhl2*). Scale bars: 200 μm. **D.** Genome browser tracks displaying chromatin accessibility (Top), peak sites, Signac-derived Peak2GeneLinks, SCENIC+-predicted eRegulonLinks (Middle), gene tracks (Bottom), and corresponding gene expression (Right panel) for TF-eRegulon-target gene pairs: (*Grhl2*+*Foxa1*)-*Sox2*. **E**. H&E staining of the adjacent slice of E12.5_3 and spatial distribution of TF target genes on the E12.5_3 section. Target genes include *Ap1s3* for *Foxa2*, *Lin7a* for *Nkx2-1*, and *Sox2* for *Grhl2*. Scale bars: 200 μm.

**Figure S8.**
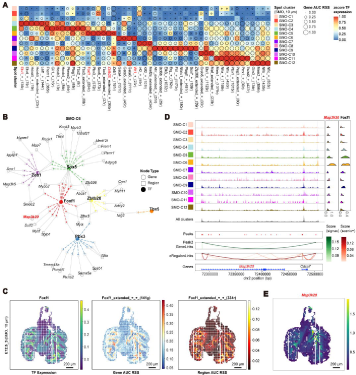
Gene regulatory networks and spatial visualization of mesenchymal clusters in early mouse lung development. **A**. Heatmap/dot plot illustrating the z-score TF expression (color scale) and cell-type specificity (RSS, size scale) of TF-eRegulons enriched in mesenchymal clusters. **B**. TF-region-gene regulatory networks for SMO-C6. Dashed lines represent TF-region regulatory relationships, while solid lines indicate region-gene regulatory interactions. **C**. Spatial mapping of spot cluster-specific TF-eRegulons, showing TF expression, Gene AUC RSS, and Region AUC RSS for SMO-C6 (Airway smooth muscle cell precursors; *Foxf1*). Scale bars: 200 μm. **D.** Genome browser tracks displaying chromatin accessibility (Top), peak sites, Signac-derived Peak2GeneLinks, SCENIC+-predicted eRegulonLinks (Middle), gene tracks (Bottom), and corresponding gene expression (Right panel) for TF-eRegulon-target gene pairs: *Map3k20*-*Foxf1*. **E**. Spatial visualization of target gene *Map3k20* for *Foxf1* on the E12.5_3 section. Scale bars: 200 μm.

**Figure S9.**
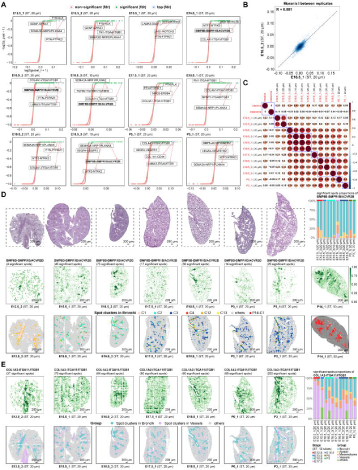
Spatial cell-cell communication during mouse lung development analyzed by SpatialDM. **A**. Scatter plots illustrating LRs enrichment in each sample using SpatialDM. The x-axis represents log2(global_I + 1), and the y-axis represents -log2(z_pval + 1). **B**. Example scatter plot showing the correlation of LR enrichment between two biological replicates at E16.5. **C**. Heatmap displaying Pearson’s correlation coefficients for shared LRs across different samples. **D**. Top: H&E staining of adjacent tissue sections. Middle: Spatial visualization of *BMP8B-BMPR1B/ACVR2B* LR spatial autocorrelation probability (1 - z_pval) in airway structures. Bottom: Spatial distribution of spot clusters within the bronchi group across different lung sections during development. Top right: Histogram showing the proportion of *BMP8B-BMPR1B/ACVR2B* interactions in different lung structure spots from E12.5 to P3. Scale bars: 200 μm. **E**. Top: H&E staining of adjacent tissue sections. Middle: Spatial visualization of *COL1A2-ITGA11/ITGB1* LR spatial autocorrelation probability (1 - z_pval) in airway and vascular structures. Bottom: Spatial distribution of spot clusters within the bronchi and vessel groups across different lung sections during development. Right panel: Histogram showing the proportion of *COL1A2-ITGA11/ITGB1* interactions in different lung structure spots from E12.5 to P3. Scale bars: 200 μm.

**Figure S10.**
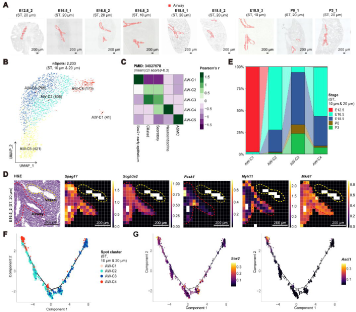
Spatial development of airway epithelial clusters in the mouse lung. **A**. Selection of lung bronchial airway spots based on manual annotation and morphological characteristics from E12.5, E16.5, E18.5, P0, and P3 lungs. **B**. Re-clustering UMAP of selected lung bronchial airway spots. AW: airway. **C**. Heatmaps displaying the distribution of scaled cell type abundances across spatial clusters for the single-cell RNA-seq datasets (PMID: 34927678). **D**. Bar plot illustrating the dynamic proportions of airway epithelial clusters during lung development. **E**. Spatial gene expression patterns of classical markers for different airway epithelial cell types, including *Spag17* (ciliated cells), *Scgb3a2* (secretory cells), *Pcsk1* (PNECs), *Myh11* (ASMCs), and *Mki67* (proliferation marker). Red dotted lines mark airways, and yellow dotted lines mark vessels. Scale bars: 200 μm. **F**. Developmental trajectory of the four airway epithelial clusters. **G**. Visualization of gene expression for classical proximal epithelial progenitor marker (*Sox2*) and PNEC marker (*Ascl1*) along the airway developmental trajectory.

**Figure S11.**
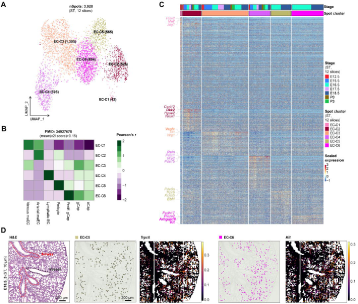
Spatial distribution of vessel-associated clusters during mouse lung development. **A**. Re-clustering UMAP visualization of vascular spots (10 μm and 20 μm-resolution ST data). **B**. Heatmaps displaying the distribution of scaled cell type abundances across vessel-associated spatial clusters for the single-cell RNA-seq datasets (PMID: 34927678). **C**. Heatmap showing normalized gene expression of differentially expressed genes (DEGs) for six spot clusters of mouse lung vascular cells. **D**. Normalized spatial gene expression patterns of cluster marker genes on ST sections, including *Trpc6* (pericyte marker) and *Kit* (general capillary cell marker). Red dotted lines on H&E staining mark airways, and black dotted lines mark large vessels adjacent to airways. Scale bars: 200 μm.

**Figure S12.**
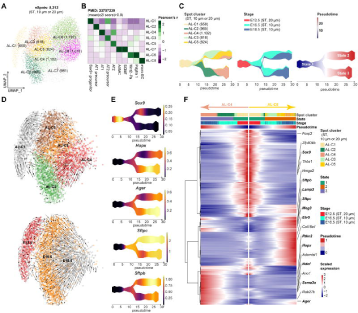
Spatial developmental trajectories of alveolar epithelial clusters during mouse lung development. **A**. Re-clustering UMAP visualization of spots in the distal airway or alveolar niche from E12.5_1 (20 μm), E12.5_2 (20 μm), E16.5_3 (10 μm), and E18.5_3 (10 μm) sections. **B**. Heatmaps displaying the distribution of scaled cell type abundances across alveolar spatial clusters for the single-cell RNA-seq dataset (PMID: 33707239). **C**. Separated developmental trajectories showing re-annotated clusters (Left), actual developmental stages (Middle), and pseudotime states (Right). **D**. RNA velocity trajectory of alveolar epithelial clusters based on the original UMAP embedding, analyzed using scVelo. **E**. Visualization of gene expression for distal/alveolar cell markers (*Sox9*, *Hopx*, *Ager*, *Sftpc*, and *Sftpb*) along the STREAM trajectory. **F**. Heatmap showing differentially expressed genes (DEGs) for alveolar epithelial clusters at different developmental states.

**Figure S13.**
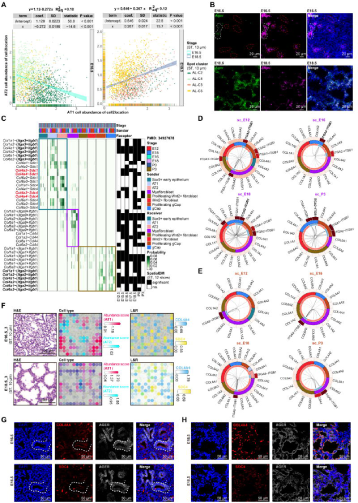
The spatial convergence of AT1 and AT2 cells mediated by collagen signaling pathways during the canalicular-saccular transition. **A**. Scatter plots showing cell2location abundance scores for AT1 and AT2 cells at E16.5 (Left) and E18.5 (Right). Linear regression was applied to assess correlations. **B**. RNA fluorescence in situ hybridization (FISH) confirming the spatial expression of *Ager* and *Sftpc* at E16.5 (Top) and E18.5 (Bottom). White dashed lines indicate *Ager*+*Sftpc*+ airways, and red dashed lines outline *Ager*-*Sftpc*+ airway tips. Scale bars: 20 μm. **C**. Heatmap showing the probability of ligand-receptor (LR) interactions from the collagen pathway within the alveolar niche. All LRs are significant in both scRNA-seq CellChat analysis and at least one ST section. **D**. Circle plot illustrating LRs involving myofibroblast cells as main receptor-expressing populations in collagen signaling pathways, derived from scRNA-seq data (PMID: 34927678). Labels are consistent with Figure 4E. **E**. Circle plot illustrating LRs involving (proliferating) gCap cells as main receptor-expressing populations in collagen signaling pathways, derived from scRNA-seq data (PMID: 34927678). Labels are consistent with Figure 4E. **F**. Example showing the spatial enrichment of *COL4A4*-*SDC4* LR in alveolar epithelial cells and regions in E16.5 and E18.5 mouse lungs. **G-H**. Immunofluorescence staining for COL4A4 and SDC4 proteins at E16.5 **(G)** and E18.5 **(H)**. COL4A4 and SDC4 co-staining with AGER was performed on two adjacent sections, respectively. Co-localization of COL4A4 and SDC4 proteins was observed at E16.5 (white dotted lines) and E18.5. Scale bars: 200 μm.

**Figure S14.**
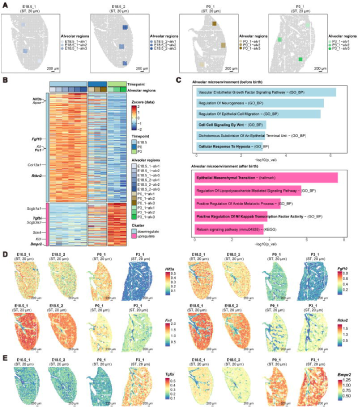
Molecular characteristics of the alveolar region before and after birth. **A**. Selected lung alveolar niche spots according to manual annotation and morphological characteristics from 20 μm-resolution E18.5, P0 and P3 data. **B**. The heatmap showing transitional alveolar niche specific genes during mouse lung alveolar maturation (E18.5, P0 and P3). **C**. Bar plots of pathway enrichment analysis for alveolar niche specific genes before (Top) or after (Bottom) birth. **D**. The spatial highlighting of normalized gene expression level for down-regulated genes after birth (*Hif3a*, *Fgf10*, *Fn1* and *Rtkn2*) across E18.5, P0 and P3. **E**. The spatial highlighting of normalized gene expression level for up-regulated genes after birth (*Tgfbi* and *Bmpr2*) across E18.5, P0 and P3.

## References

[1] Morrisey EE, Hogan BL. Preparing for the first breath: genetic and cellular mechanisms in lung development. Dev Cell 2010;18:8–23.

[2] Herriges M, Morrisey EE. Lung development: orchestrating the generation and regeneration of a complex organ. Development 2014;141:502–13.

[3] Aros CJ, Pantoja CJ, Gomperts BN. Wnt signaling in lung development, regeneration, and disease progression. Commun Biol 2021;4:601.

[4] Saito A, Horie M, Nagase T. TGF-β Signaling in Lung Health and Disease. Int J Mol Sci 2018;19.

[5] Zepp JA, Morrisey EE. Cellular crosstalk in the development and regeneration of the respiratory system. Nat Rev Mol Cell Biol 2019;20:551–66.

[6] Schittny JC. Development of the lung. Cell Tissue Res 2017;367:427–44.

[7] Mullassery D, Smith NP. Lung development. Semin Pediatr Surg 2015;24:152–5.

[8] Treutlein B, Brownfield DG, Wu AR, Neff NF, Mantalas GL, Espinoza FH, et al. Reconstructing lineage hierarchies of the distal lung epithelium using single-cell RNA-seq. Nature 2014;509:371–5.

[9] Zepp JA, Zacharias WJ, Frank DB, Cavanaugh CA, Zhou S, Morley MP, et al. Distinct Mesenchymal Lineages and Niches Promote Epithelial Self-Renewal and Myofibrogenesis in the Lung. Cell 2017;170:1134–48.e10.

[10] Negretti NM, Plosa EJ, Benjamin JT, Schuler BA, Habermann AC, Jetter CS, et al. A single-cell atlas of mouse lung development. Development 2021;148.

[11] Cohen M, Giladi A, Gorki AD, Solodkin DG, Zada M, Hladik A, et al. Lung Single-Cell Signaling Interaction Map Reveals Basophil Role in Macrophage Imprinting. Cell 2018;175:1031–44.e18.

[12] Travaglini KJ, Nabhan AN, Penland L, Sinha R, Gillich A, Sit RV, et al. A molecular cell atlas of the human lung from single-cell RNA sequencing. Nature 2020;587:619–25.

[13] Zepp JA, Morley MP, Loebel C, Kremp MM, Chaudhry FN, Basil MC, et al. Genomic, epigenomic, and biophysical cues controlling the emergence of the lung alveolus. Science 2021;371.

[14] Basil MC, Cardenas-Diaz FL, Kathiriya JJ, Morley MP, Carl J, Brumwell AN, et al. Human distal airways contain a multipotent secretory cell that can regenerate alveoli. Nature 2022;604:120–6.

[15] Price D, Graham L, Moss J, Armit C, Baldock RA. Kaufman’s Atlas of Mouse Development Supplement 2015.

[16] Rao A, Barkley D, França GS, Yanai I. Exploring tissue architecture using spatial transcriptomics. Nature 2021;596:211–20.

[17] Bressan D, Battistoni G, Hannon GJ. The dawn of spatial omics. Science 2023;381:eabq4964.

[18] Liu Y, Yang M, Deng Y, Su G, Enninful A, Guo CC, et al. High-Spatial-Resolution Multi-Omics Sequencing via Deterministic Barcoding in Tissue. Cell 2020;183:1665–81.e18.

[19] Kleshchevnikov V, Shmatko A, Dann E, Aivazidis A, King HW, Li T, et al. Cell2location maps fine-grained cell types in spatial transcriptomics. Nat Biotechnol 2022;40:661–71.

[20] Jiang F, Zhou X, Qian Y, Zhu M, Wang L, Li Z, et al. Simultaneous profiling of spatial gene expression and chromatin accessibility during mouse brain development. Nat Methods 2023;20:1048–57.

[21] Lin S, Perl AK, Shannon JM. Erm/thyroid transcription factor 1 interactions modulate surfactant protein C transcription. J Biol Chem 2006;281:16716–26.

[22] Bravo González-Blas C, De Winter S, Hulselmans G, Hecker N, Matetovici I, Christiaens V, et al. SCENIC+: single-cell multiomic inference of enhancers and gene regulatory networks. Nat Methods 2023;20:1355–67.

[23] Wan H, Dingle S, Xu Y, Besnard V, Kaestner KH, Ang SL, et al. Compensatory roles of Foxa1 and Foxa2 during lung morphogenesis. J Biol Chem 2005;280:13809–16.

[24] Park HL, Bai C, Platt KA, Matise MP, Beeghly A, Hui CC, et al. Mouse Gli1 mutants are viable but have defects in SHH signaling in combination with a Gli2 mutation. Development 2000;127:1593–605.

[25] Shaw-White JR, Bruno MD, Whitsett JA. GATA-6 activates transcription of thyroid transcription factor-1. J Biol Chem 1999;274:2658–64.

[26] He P, Lim K, Sun D, Pett JP, Jeng Q, Polanski K, et al. A human fetal lung cell atlas uncovers proximal-distal gradients of differentiation and key regulators of epithelial fates. Cell 2022;185:4841–60.e25.

[27] Soler Artigas M, Loth DW, Wain LV, Gharib SA, Obeidat M, Tang W, et al. Genome-wide association and large-scale follow up identifies 16 new loci influencing lung function. Nat Genet 2011;43:1082–90.

[28] Pan L, Hoffmeister P, Turkiewicz A, Huynh NND, Große-Berkenbusch A, Knippschild U, et al. Transcription Factor RBPJL Is Able to Repress Notch Target Gene Expression but Is Non-Responsive to Notch Activation. Cancers (Basel) 2021;13.

[29] Mendelson CR. Role of transcription factors in fetal lung development and surfactant protein gene expression. Annu Rev Physiol 2000;62:875–915.

[30] Li C, Cai J, Pan Q, Minoo P. Two functionally distinct forms of NKX2.1 protein are expressed in the pulmonary epithelium. Biochem Biophys Res Commun 2000;270:462–8.

[31] Gruel N, Fuhrmann L, Lodillinsky C, Benhamo V, Mariani O, Cédenot A, et al. LIN7A is a major determinant of cell-polarity defects in breast carcinomas. Breast Cancer Res 2016;18:23.

[32] He S, Li Y, Shi X, Wang L, Cai D, Zhou J, et al. DNA methylation landscape reveals LIN7A as a decitabine-responsive marker in patients with t(8;21) acute myeloid leukemia. Clin Epigenetics 2023;15:37.

[33] Hu X, Li Y, Kong D, Hu L, Liu D, Wu J. Long noncoding RNA CASC9 promotes LIN7A expression via miR-758-3p to facilitate the malignancy of ovarian cancer. J Cell Physiol 2019;234:10800–8.

[34] Luo C, Yin D, Zhan H, Borjigin U, Li C, Zhou Z, et al. microRNA-501-3p suppresses metastasis and progression of hepatocellular carcinoma through targeting LIN7A. Cell Death Dis 2018;9:535.

[35] Kersbergen A, Best SA, Dworkin S, Ah-Cann C, de Vries ME, Asselin-Labat M-L, et al. Lung morphogenesis is orchestrated through Grainyhead-like 2 (Grhl2) transcriptional programs. Developmental Biology 2018;443:1–9.

[36] Wang H, St Julien KR, Stevenson DK, Hoffmann TJ, Witte JS, Lazzeroni LC, et al. A genome-wide association study (GWAS) for bronchopulmonary dysplasia. Pediatrics 2013;132:290–7.

[37] Nawijn MC, Hackett TL, Postma DS, van Oosterhout AJ, Heijink IH. E-cadherin: gatekeeper of airway mucosa and allergic sensitization. Trends Immunol 2011;32:248–55.

[38] Que J, Luo X, Schwartz RJ, Hogan BL. Multiple roles for Sox2 in the developing and adult mouse trachea. Development 2009;136:1899–907.

[39] Boucherat O, Landry-Truchon K, Aoidi R, Houde N, Nadeau V, Charron J, et al. Lung development requires an active ERK/MAPK pathway in the lung mesenchyme. Dev Dyn 2017;246:72–82.

[40] Polyak K, Kalluri R. The role of the microenvironment in mammary gland development and cancer. Cold Spring Harb Perspect Biol 2010;2:a003244.

[41] De Belly H, Paluch EK, Chalut KJ. Interplay between mechanics and signalling in regulating cell fate. Nat Rev Mol Cell Biol 2022;23:465–80.

[42] Quach H, Farrell S, Wu MJM, Kanagarajah K, Leung JW, Xu X, et al. Early human fetal lung atlas reveals the temporal dynamics of epithelial cell plasticity. Nat Commun 2024;15:5898.

[43] Li Z, Wang T, Liu P, Huang Y. SpatialDM for rapid identification of spatially co-expressed ligand-receptor and revealing cell-cell communication patterns. Nat Commun 2023;14:3995.

[44] Yang L, Zhou F, Zheng D, Wang D, Li X, Zhao C, et al. FGF/FGFR signaling: From lung development to respiratory diseases. Cytokine Growth Factor Rev 2021;62:94–104.

[45] Zong D, Ouyang R, Li J, Chen Y, Chen P. Notch signaling in lung diseases: focus on Notch1 and Notch3. Ther Adv Respir Dis 2016;10:468–84.

[46] Caldeira I, Fernandes-Silva H, Machado-Costa D, Correia-Pinto J, Moura RS. Developmental Pathways Underlying Lung Development and Congenital Lung Disorders. Cells 2021;10.

[47] Weng T, Liu L. The role of pleiotrophin and beta-catenin in fetal lung development. Respir Res 2010;11:80.

[48] Ghosh S, Paez-Cortez JR, Boppidi K, Vasconcelos M, Roy M, Cardoso W, et al. Activation dynamics and signaling properties of Notch3 receptor in the developing pulmonary artery. J Biol Chem 2011;286:22678–87.

[49] Warburton D, Bellusci S, De Langhe S, Del Moral PM, Fleury V, Mailleux A, et al. Molecular mechanisms of early lung specification and branching morphogenesis. Pediatr Res 2005;57:26r–37r.

[50] Vacca M, Leslie J, Virtue S, Lam BYH, Govaere O, Tiniakos D, et al. Bone morphogenetic protein 8B promotes the progression of non-alcoholic steatohepatitis. Nat Metab 2020;2:514–31.

[51] Whittle AJ, Carobbio S, Martins L, Slawik M, Hondares E, Vázquez MJ, et al. BMP8B increases brown adipose tissue thermogenesis through both central and peripheral actions. Cell 2012;149:871–85.

[52] Zhao GQ, Deng K, Labosky PA, Liaw L, Hogan BL. The gene encoding bone morphogenetic protein 8B is required for the initiation and maintenance of spermatogenesis in the mouse. Genes Dev 1996;10:1657–69.

[53] Song H, Yao E, Lin C, Gacayan R, Chen MH, Chuang PT. Functional characterization of pulmonary neuroendocrine cells in lung development, injury, and tumorigenesis. Proc Natl Acad Sci U S A 2012;109:17531–6.

[54] Sui P, Wiesner DL, Xu J, Zhang Y, Lee J, Van Dyken S, et al. Pulmonary neuroendocrine cells amplify allergic asthma responses. Science 2018;360.

[55] Branchfield K, Nantie L, Verheyden JM, Sui P, Wienhold MD, Sun X. Pulmonary neuroendocrine cells function as airway sensors to control lung immune response. Science 2016;351:707–10.

[56] Kuo CS, Krasnow MA. Formation of a Neurosensory Organ by Epithelial Cell Slithering. Cell 2015;163:394–405.

[57] Noguchi M, Sumiyama K, Morimoto M. Directed Migration of Pulmonary Neuroendocrine Cells toward Airway Branches Organizes the Stereotypic Location of Neuroepithelial Bodies. Cell Rep 2015;13:2679–86.

[58] Chen H, Albergante L, Hsu JY, Lareau CA, Lo Bosco G, Guan J, et al. Single-cell trajectories reconstruction, exploration and mapping of omics data with STREAM. Nat Commun 2019;10:1903.

[59] Frank DB, Penkala IJ, Zepp JA, Sivakumar A, Linares-Saldana R, Zacharias WJ, et al. Early lineage specification defines alveolar epithelial ontogeny in the murine lung. Proc Natl Acad Sci U S A 2019;116:4362–71.

[60] Desai TJ, Brownfield DG, Krasnow MA. Alveolar progenitor and stem cells in lung development, renewal and cancer. Nature 2014;507:190–4.

[61] Rawlins EL, Clark CP, Xue Y, Hogan BL. The Id2+ distal tip lung epithelium contains individual multipotent embryonic progenitor cells. Development 2009;136:3741–5.

[62] Bergen V, Lange M, Peidli S, Wolf FA, Theis FJ. Generalizing RNA velocity to transient cell states through dynamical modeling. Nat Biotechnol 2020;38:1408–14.

[63] Frank DB, Peng T, Zepp JA, Snitow M, Vincent TL, Penkala IJ, et al. Emergence of a Wave of Wnt Signaling that Regulates Lung Alveologenesis by Controlling Epithelial Self-Renewal and Differentiation. Cell Rep 2016;17:2312–25.

[64] Barkauskas CE, Cronce MJ, Rackley CR, Bowie EJ, Keene DR, Stripp BR, et al. Type 2 alveolar cells are stem cells in adult lung. J Clin Invest 2013;123:3025–36.

[65] Sainio A, Järveläinen H. Extracellular matrix-cell interactions: Focus on therapeutic applications. Cell Signal 2020;66:109487.

[66] Schmidt S, Friedl P. Interstitial cell migration: integrin-dependent and alternative adhesion mechanisms. Cell Tissue Res 2010;339:83–92.

[67] Shiraishi K, Shah PP, Morley MP, Loebel C, Santini GT, Katzen J, et al. Biophysical forces mediated by respiration maintain lung alveolar epithelial cell fate. Cell 2023;186:1478–92.e15.

[68] Gopal S, Arokiasamy S, Pataki C, Whiteford JR, Couchman JR. Syndecan receptors: pericellular regulators in development and inflammatory disease. Open Biol 2021;11:200377.

[69] Swarr DT, Morrisey EE. Lung endoderm morphogenesis: gasping for form and function. Annu Rev Cell Dev Biol 2015;31:553–73.

[70] Huang Y, Kapere Ochieng J, Kempen MB, Munck AB, Swagemakers S, van Ijcken W, et al. Hypoxia inducible factor 3α plays a critical role in alveolarization and distal epithelial cell differentiation during mouse lung development. PLoS One 2013;8:e57695.

[71] Jiang J, Zheng Z, Chen S, Liu J, Jia J, Huang Y, et al. Hypoxia inducible factor (HIF) 3α prevents COPD by inhibiting alveolar epithelial cell ferroptosis via the HIF-3α-GPx4 axis. Theranostics 2024;14:5512–27.

[72] Liu M, Iosef C, Rao S, Domingo-Gonzalez R, Fu S, Snider P, et al. Transforming Growth Factor-induced Protein Promotes NF-κB-mediated Angiogenesis during Postnatal Lung Development. Am J Respir Cell Mol Biol 2021;64:318–30.

[73] Chénais B. Transposable elements and human cancer: a causal relationship? Biochim Biophys Acta 2013;1835:28–35.

[74] Wells JN, Feschotte C. A Field Guide to Eukaryotic Transposable Elements. Annu Rev Genet 2020;54:539–61.

[75] Yu H, Chen M, Hu Y, Ou S, Yu X, Liang S, et al. Dynamic reprogramming of H3K9me3 at hominoid-specific retrotransposons during human preimplantation development. Cell Stem Cell 2022;29:1031–50.e12.

[76] Herriges JC, Verheyden JM, Zhang Z, Sui P, Zhang Y, Anderson MJ, et al. FGF-Regulated ETV Transcription Factors Control FGF-SHH Feedback Loop in Lung Branching. Dev Cell 2015;35:322–32.

[77] Chung MI, Bujnis M, Barkauskas CE, Kobayashi Y, Hogan BLM. Niche-mediated BMP/SMAD signaling regulates lung alveolar stem cell proliferation and differentiation. Development 2018;145.

