## Supplementary material for "Spatial Transcriptomics Unveils the Blueprint of Mammalian Lung Development": Materials and methods

**Animals**

C57BL/6 background mice were used in this study. Timed-pregnant C57BL/6 females were utilized for all embryonic time points, with pregnancy stages confirmed by crown-rump length measurements. Embryonic day (E) 0.5 was designated as noon on the day a vaginal plug was observed, and postnatal day (P) 0 was defined as the day of birth. All mice were housed in the Guangzhou National Laboratory animal facility under controlled conditions, including a 12-hour light-dark cycle, constant temperature, and humidity, and were maintained in a healthy state throughout the study.

**Sample collecting**

Given the significant role of sex and sex hormones in lung development under both physiological and pathophysiological conditions, this study exclusively utilized male mice unless otherwise specified [1-4]. The sex of postnatal mice was determined visually, while PCR was employed for sex determination in embryos. PCR was performed using the following primers: Sry-F (5'-ATGAATGCATTTATGGTGTGGTCCCGTGGTGAGAG-3') and Sry-R (5'-GAGTACAGGTGTGCAGCTCTACTCCAGTCTTGCCT-3'). Fresh frozen lung tissues were collected at E12.5, E15.5, E16.5, E17.5, E18.5, P0, P3, and P14 for spatial transcriptomics (ST) experiments. E12.5 tissues were used for modified MISAR-seq (10 μm resolution) and DBiT-seq (20 μm resolution), while P14 tissues were exclusively used for 500 nm-resolution Stereo-seq library construction. Notably, for P14 samples, blood was removed from the eyes after anesthesia, followed by lung perfusion with 50% OCT (catalogue no.A4583, Torrance, CA) in PBS (v/v) and embedding in OCT. The embedded tissues were immediately stored at -80 °C until sectioning for subsequent experiments.

**Microfluidic device obtaination**

Fifty 20 μm-wide and ninety-six 10/20 μm-wide polydimethylsiloxane (PDMS) microfluidic chips, along with their matched clamps, custom unembedded slide glass (84.9 × 46.54 × 1 mm), and corresponding storage boxes, were procured from ZhongXinQiHeng (Suzhou Chip Scientific Instrument Co., Ltd.). The channel interval thickness of the chips was approximately 10 μm for 10 μm-wide chips and 20 μm for 20 μm-wide chips, respectively. Custom poly-L-lysine (catalogue no.P8920-500ML, St. Louis, Missouri) coated glass slides were prepared following the manufacturer's instructions.

**Tissue cryosection**

Prior to sectioning, frozen embedded tissues and slides were equilibrated in the cryostat for at least 30 minutes to reach the cryostat temperature (~ -20°C). Tissue sections of 10 μm thickness were prepared for the 20 μm-wide microfluidic channel device, while 8 μm sections were used for the 10 μm-wide device. Sections were carefully positioned at the center of pre-marked commercial or custom glass slides, depending on tissue size. During the sectioning process, approximately ten tissue sections were collected into 1.5 mL centrifuge tubes for subsequent RNA quality assessment.

**RNA quality identification**

RNA was extracted using the Quick-RNA™ MicroPrep Kit (catalogue no.R1050, Irvine, CA) following the manufacturer's protocol. RNA quality was assessed through agarose gel electrophoresis. Only sections meeting the RNA quality criteria were selected for subsequent spatial transcriptome experiments.

**H&E staining**

To assess histological morphology, adjacent tissue sections from the spatial experiment were subjected to hematoxylin and eosin (H&E) staining using a Hematoxylin and Eosin Staining Kit (catalogue no.C0105S, Shanghai, China) . Briefly, air-dried slides were washed with 1× PBS and fixed with 10% neutral-buffered formalin (NBF) for 10 minutes. After rinsing with distilled water, tissues were stained with hematoxylin solution for 2–4 minutes and immediately rinsed with distilled water. The sections were then treated with hydrochloric acid differentiation solution for approximately 10 seconds. Following another wash, the tissues were counterstained with eosin for about 8 minutes. Excess eosin was removed using 75% ethanol after a final rinse with distilled water. Slides were air-dried and mounted with coverslips. Stained tissues were imaged using the EVOS™ M7000 Imaging System (catalog no. AMF7000, Carlsbad, CA) at 200× magnification.

**Transposome embedding for spatial transcriptome library construction**

Unmodified Tn5 transposase was purchased from Novoprotein (catalog no. M045-01B, Suzhou, China). The transposome complex was assembled according to the manufacturer's instructions using TN5_Read1_RNA and TN5_ME_RNA adapters (sequences provided in Table S1).

**Spatial transcriptome library construction with modified DBiT-seq**

Spatial RNA-seq library construction was performed based on the principle of DBiT-seq [5] with minor modifications.

Briefly, tissue sections were initially cleaned with PBS-RI, fixed with 4% formaldehyde (FA), and permeabilized with 0.05% Triton X-100 for 20 minutes at room temperature. Reverse transcription was then performed by incubating the sections with RT primer at room temperature for 30 minutes, followed by 42°C for 90 minutes. Spatial barcoding of mRNAs was achieved using two rounds of PDMS microfluidic chips. Annealed barcode A or B mixtures, blocking A or B mixtures, and 1× NEB buffer 3.1 were sequentially added to the corresponding chip inlets and flowed across the tissue surface through the channels via vacuum. Reverse crosslinking was performed using lysis solution containing proteinase K, and the lysate was collected and stored at -80°C until further processing.

Lysate purification was carried out using the DNA Clean and Concentrator kit(catalog no. D4013, Irvine, CA) according to the manufacturer’s instructions. cDNA was captured using MyOne™ Streptavidin C1 beads (catalog no. 65001, Carlsbad, CA). After washing with B&W and STE solutions, the beads were resuspended in template switch mix containing TSO oligos. Following template switching, the beads were washed with STE and nuclease-free water (NF-H_2_O), resuspended in PCR mix with cDNA_ampli_R and RNA_PCR_primer, and amplified for 10–13 cycles. Amplified products were purified using 0.8× SPRIselect beads (catalog no. 65001, Brea, CA). cDNA quantity was quantified using a Qubit fluorometer (Thermo Fisher Scientific). For each sample, 50 ng of cDNA was fragmented in a 20 μL tagmentation reaction with custom Tn5 transposase, and the tagmented products were purified using 2× SPRIselect beads. The 3' end products were amplified with Illumina Nextera N5 and N7 adapters and purified with 0.7× SPRIselect beads. After quality assessment, libraries were sequenced on the NovaSeq 6000 platform with 150-bp paired-end reads. Oligonucleotide sequences used in this protocol are listed in Table S1.

**Stereo-seq**

P14 lung tissue blocks were prepared and sectioned as described above. Stereo-seq libraries were constructed following previously established protocols [6]. Briefly, P14 lung samples were cryosectioned at 10 μm thickness after equilibration in the cryostat for 30 minutes. The first section was mounted onto the Stereo-seq chip surface, and the subsequent four sections were used to determine the optimal permeabilization time, which was determined to be 6 minutes. Library construction was performed according to standard procedures, and paired-end 100 bp (PE100) sequencing was conducted.

**Spatial assay for ATAC and RNA-sequencing with modified MISAR-seq**

Spatial ATAC-seq library construction was performed based on the principle of MISAR-seq [7] with minor modifications. Briefly, fresh E12.5 mouse lung sections were subjected to fixation, permeabilization, and transposition. Custom Tn5 transposase, provided by Novoprotein, was used for transposition, along with adapters (ATAC_Phosphorylated_Read2, ATAC_Read1, and ATAC_Blocked_ME_Comp), as listed in Table S1. Reverse transcription, ligation, and lysis reactions were carried out following the same procedures as described for spatial transcriptome library construction. MyOne™ Streptavidin C1 beads were used to separate cDNA and genomic DNA, and cDNA library construction was performed as outlined above.

For ATAC library preparation, genomic DNA retained in the supernatant after C1 bead separation was purified using the Zymo DNA Clean & Concentrator-5 kit and eluted in 21 μL of nuclease-free water (NF-H_2_O). The eluted product was mixed with PCR solution (2.5 μL of 10 μM Illumina Nextera N5 adapters, 2.5 μL of 10 μM Illumina Nextera N7 adapters, and 25 μL of 2× NEBNext Master Mix). PCR amplification was performed using the following program: 72 °C for 5 minutes, 98 °C for 30 seconds, followed by 9 cycles of 98 °C for 10 seconds, 65 °C for 30 seconds, and 72 °C for 1 minute, with a final extension at 72 °C for 5 minutes. PCR products were purified using 1.2× SPRIselect beads and resuspended in 20 μL of RNase-free water for sequencing.

**Immunofluorescence staining**

P14 and adult male mice were anesthetized with isoflurane and perfused with PBS through the right ventricle. Lungs were inflated with 10% neutral-buffered formalin (NBF) and fixed in 10% NBF at 4°C overnight. Fixed lungs were cryoprotected in PBS containing 20% sucrose and 33% OCT at 4°C overnight, followed by embedding in OCT. For embryonic time points, lungs were dissected from timed-pregnant mice and fixed in 10% NBF at 4°C overnight. Tissues were then gradiently dehydrated in 10%, 20%, and 30% sucrose solutions in PBS, embedded in OCT, and stored at -80°C until further processing.

Immunofluorescence staining was performed using the following primary antibodies: NKX2.1 (catalog no. WRAB-1231, Cincinnati, OH; 1:200), ACTA2 (catalog no. MA1-06110, Waltham, MA; 1:250), AGER (catalog no. MAB1179-SP, Minneapolis, MN; 1:200), SFTPC (catalog no. ABC99, St. Louis, MO; 1:200), SDC4 (catalog no. 11820-1-AP, Wuhan, China; 1:100), and COL4A4 (catalog no. 19674-1-AP, Wuhan, China; 1:100). Alexa Fluor secondary antibodies included anti-mouse 488 (catalog no. 4408S, Beverly, MA), anti-rat 555 (catalog no. 4417S, Beverly, MA), anti-rabbit 546 (catalog no. 4417S, Beverly, MA), and anti-rabbit 647 (catalog no. ab150075, Cambridge, UK). Images were acquired using a laser confocal microscope (LSM 980, Zeiss) with 10×, 20×, or 40× water-immersion objectives at random fields.

**RNA Fluorescence In Situ Hybridization (FISH)**

Target-specific oligonucleotide probes (20–50 nt) conjugated with fluorophores (e.g., Cy3, Alexa 488) were designed and synthesized by Zoonbio Biotechnology Co., Ltd., and frozen tissue sections (10 μm thickness) from E16.5 and E18.5 lungs were prepared for RNA FISH experiments. Briefly, after washing with PBS, tissue sections were treated with proteinase K working solution at 37 °C for 20 minutes. Sections were then incubated with prehybridization solution at 37 °C for 1 hour to block nonspecific binding. Subsequently, diluted probes were applied to the sections and hybridized overnight at 42 °C in a humidified chamber. Following hybridization, sections were washed with gradient SSC buffers to remove unbound probes. Nuclei were counterstained with DAPI, and slides were mounted using an anti-fade mounting medium to preserve fluorescence. Imaging was performed using a Nikon inverted fluorescence microscope, and images were acquired for downstream analysis.

**Spatial RNA data pre-processing and quality control**

To get gene-spot count matrices, we first trimmed adapters of sequence reads using Cutadapt (v4.2) [8], then used UMI-tools (v1.1.2) [9] to debarcode. For the final counting step, we utilized zUMIs (v2.9.7d) [10] to get UMIs for each spot based on the reference genome (GRCm38) and the corresponding gtf (Ensemble release 102).

To ensure the validity of the spots captured by sequencing, we discarded any spot which was not covered on the tissue when mapping to the FIX image. In order to filter low quality spots, we dropped spots with fewer than 500 genes detected or 1,000 UMIs before later analyses. Additionally, spots with high level of blood associated genes were excluded, and this cutoff varied within specific slice. All statistical analysis in violin plots and box plots were produced using `wilcox.test(alternative=“two.sided)` of R package `stats`.

**Stereo-seq data pre-processing and quality control**

The raw fastq data for Stereo-seq analysis was processed with the publicly available SAW pipeline (version v5.4.0) (https://github.com/STOmics/SAW), mapping to the reference genome (GRCm38) and the corresponding gtf (Ensemble release 102). After that we proceeded the gene expression profile matrices within non-overlapping bins of 20 x 20, 40 x 40, 50 x 50, 100 x 100 and 200 x 200 DNB, respectively. We selected bin100 (50um) resolution for further analysis, and filtered low quality spots followed the protocol previously described.

**Spatial RNA data normalizing, integrating, clustering and annotation**

We normalized each raw gene-spot matrices individually by SCTransform [11] (vst.flavor="v2", method="glmGamPoi")` using Seurat (v4.3.0) [12]. As there were no batch effects on different replicates, we chose highly variable genes (HVGs) across samples through `SelectIntegrationFeatures()` to re-normalize each sample individually through the same method, and combined all samples via `merge()`. In order to reduce dimensionality of the data, we performed `RunPCA()` to select the suitable number of PC which had a cumulative variance contribution ratio higher than 80%. And then we used `FindNeighbors()` to construct a KNN graph based on selected PCs. For unsupervised clustering of spots, we used the Louvain algorithm via `FindClusters()` with resolution from 0.1 to 2.0, and projected data to a 2D plane via `RunUMAP()` for visualization. Considering noise spots and rare cell types, we further performed subclustering on each cluster with resolution from 1.0 to 2.0, clusters with significant biological features were remained while noise clusters were excluded in the later analysis.

To annotate each cluster, we first collected a set of classical lung cell type markers from the published database [13, 14], including their classical cell signatures and abundance. We first divided clusters into 5 main groups: Alveolar epithelium, Bronchial epithelium, Vascular endothelium, Vascular fibroblast, Mesenchyme and Others. Immune cells were not excluded during analysis but were not resolved as distinct clusters due to low abundance in prenatal stages and technical sensitivity limits.Then we re-annotated each main group through high resolution clustering, cell type features enrichment and spatial distribution. Considering the resolution of ST data is not exactly single-cell, we annotated each cluster using gene sets via `AddModuleScore()` rather than individual genes. To verify the rationality of annotation, pearson correlations were calculated between each cluster via `cor()` in R, the result showed high biological correalations within sub-clusters. In order to find differentially expressed genes (DEGs) for each annotated cluster, we performed Wilcoxon Rank Sum test for each gene across clusters via `FindAllMarkers(logfc.threshold = 0.3, min.pct = 0.15)`, the DEGs of each cluster corresponded well to its biological features. In total, we annotate 24 clusters, each with its own unique biological features and corresponding spatial distribution.

**Spatial deconvolution of Spatial RNA data**

To get exact cell type proportions of each spot, we used both spacexr (v2.2.0) [15] and Cell2location(v0.1.3) [16] to deconvolute each spatial slice. To get comprehensive single-cell reference of mouse lung development, we downloaded the public scRNA-seq dataset [17] and re-annotated the dataset by timepoint including E12, E15, E16, E18, P0 and P3 seperately. To deconvolute spatial spots rationally, we prepared single-cell reference and spatial datasets correspondingly according to the timepoint.

Robust Cell Type Decomposition (RCTD, spacexr) can learn cell type profiles from scRNA-seq dataset to label spatial transcriptomics spots as cell types. Additionally, RCTD has a platform effect normalization step, which normalizes the scRNA-seq cell type profiles to match the platform effects of the spatial transcriptomics dataset. We used raw count matrices as input for both single-cell reference and spatial datasets, and ran RCTD in `full mode` with default paremeters. We further processed output results to visualize cell proportions of each spot in pie chart via `geom_pie()`.

Cell2location is a principled Bayesian model that can estimate which combination of cell types in which cell abundance could have given the mRNA counts in the spatial data while modelling technical effects. Similar to RCTD, we used raw count matrices to estimate absolute spatial abundance of cell types following the official tutorial of Cell2location, additionally, we set `N_cells_per_location=4` according to spatial distribution of DBIT-seq spot, and other parameters of Cell2location were remained default.

**Spatial ATAC data pre-processing and quality control**

We first trimmed adapters of sequence reads using Cutadapt (v4.2) [8] and trimmomatic (v0.39) [18], then used a custom python script to debarcode. Fragment files and feature barcode matrices were generated using scATAC-pro (v1.3.0) [19] based on the reference genome (GRCm38).

We filtered out valid spots the same as ST RNA data pre-processing and the filtered matrices were then read into ArchR (v1.0.2) as a tile matrix using createArrowFiles() and ArchRProject(). Furthermore, low quality spots with lower than 4 TSS score and fewe than 1000 Fragments were dropped before downstream analysis. Finally we remained spots both valid in ST RNA and ATAC, and generated fragment files into Signac (v1.13.0) using CreateFragmentObject() , FeatureMatrix() and CreateChromatinAssay(). Peak sites and Peak2GeneLinks were then predicted through RegionStats() and LinkPeaks().

**Spatial ATAC data normalizing, integrating, clustering and coembedding with RNA**

We firstly combined peaks of two E12.5 replicates based on tutorial of Signac (v1.13.0), and then re-created Signac-object via CreateFragmentObject() , FeatureMatrix() and CreateChromatinAssay() using unified peaks. `RunTFIDF()` was used to normalize the merged spot-peak matrix, and `FindTopFeatures()` was applied to chose highly variable peaks (HVPs). In order to reduce dimensionality of the data, we performed `RunSVD()` based on HVPs called previously. And then we used `FindNeighbors()` to construct a KNN graph based on selected LSIs. Finally `FindClusters()` with resolution from 0.1 to 2.0 was applied to do unsupervised clustering, and `RunUMAP()` was applied to project data to a 2D plane for visualization.

To project both RNA and ATAC into a single embedding, we used `FindMultiModalNeighbors()` in Seurat (v.4.3.0) to construct a weighted nearest neighbor

(WNN) graph by combining dimensional reductions computed for both modalities. To identify clusters based on co-embedding, we further generate a shared nearest neighbor graph via `FindNeighbors()` and do unsupervised clustering via `FindClusters()`. For two-dimensional visualization of the multi-omics data, UMAP space was projected via `RunUMAP()` based on co-embedding.

**Transcription factor (TF) analysis of E12.5 lung ATAC-seq data**

The SCENIC+ (v1.0.0) [20, 21] pipeline was applied to predict transcription factors (TF) with related target genes and quantify both TF activity and region enrichment at E12.5, using combined stRNA-seq and stATAC-seq data.

We firstly used pycisTopic to detect candidate enhancer regions based on cell type-specific different chromatin accessibility regions (DARs) and topics, and then potential TF binding sites (TFBSs) in candidate enhancers were identified by Motif enrichment analysis. To construct the TF-gene regulatory network, we utilized GRNBoost2 to quantify the importance of TFs and candidate enhancers to target genes and infered the direction of regulation (activation/inhibition) using linear correlations. We then visualizaed the TF-gene regulatory network via `plot_networkx()` to identify the key regulators. Finally we combined Motif enrichment analysis results with GRNBoost2 inference to recover the optimal TF of each group of motifs and generate eRegulon (enhancer-driven regulon).

In order to screen out significant eRegulons, we firstly filtered out activated eRegulons with rho > 0.4, and then selected eRegulons that both TF and target gene should be significant at 3 levels: RNA, ATAC and spatial.

**Peak-to-gene linkage analysis**

In order to identify putative regulatory relationships between peaks (ATAC) and genes (RNA), we calculated correlations of peak accessibility and gene expression using ArchR (v1.0.2)’s addPeak2GeneLinks(reducedDims = "LSI_Combined", dimsToUse = 1:60). The function firstly imputed the gene expression values based on the anchor-weight matrix. Next, the function generated cell groups via a k-nearest neighbor algorithm to reduce noise and enhance robustness, groups with higher than 80% overlap with any other were removed. Finally, we visualized the selected correlation between peaks and genes through plotPeak2GeneHeatmap(corCutOff = 0.3). To mark selected peaks and genes, we further predicted motifs of each TF through Signac (v1.13.0)’s AddMotifs() with database of JASPAR2024. Finally we generated the heatmap showing significant peak-to-gene linkages at E12.5 as shown in Fig2 A, we then marked peak-to-gene linkages ovarlapped with Foxf1, Nkx2-1 and Foxa2’s motif regions using ComplexHeatmap (v2.20.0)’s Heatmap() and rowAnnotation().

**Moran’s I for spatial auto-correlation**

Moran's I is widely used to evaluate spatial auto-correlation for individual feature, in order to evaluate spatial auto-correlation between two features, we adapted the extended Moran's I [22].

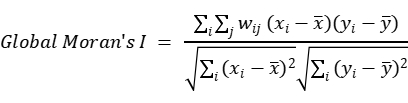

where x_i_ and y_i_ denotes normalized continuous variables of spot_i_ and spot_j_ , and w_ij_ denotes the spatial weight of spot_i_ and spot_j_ , which is calculated based on RBF kernel with an element-wise normalization.

In this study, we applied Moran’s I to illustrate the dynamic spatial auto-correlation between AT1 and AT2 epithelial cells during development.

**Linear regression between AT1 and AT2 alveolar epithelial cells**

To validate the spatial gatheration of AT1 and AT2 alveolar epithelial cells during development, we used ggplot2 (v3.4.4) to generate the scatter plot of alveolar epithelial cells by E16.5 and E18.5, and then applied liner model to fit RNA expression of each spot.

**Airway trajectory inference analysis**

To derive the pseudotime time of airway cells based on transcriptome similarity, we used monocle2 (v2.26.0) [23] to project data into a reduced dimensional space while constructing a principal tree which passes through the middle of the data simultaneously, which is based on reversed graph embedding (RGE). We applied `reduceDimension(residualModelFormulaStr = "~nCount_Spatial + nFeature_Spatial")` to remove technical effects before dimensionality reduction and clustering. For detecting key driver genes that influence cell trajectory, we screened the differentially expressed genes with `avg_log2FC > 0.35 and p_val_adj< 0.03`, and then generated the pseudotime heatmap using pheatmap (V1.0.12).

**Alveolar trajectory inference analysis**

To reconstruct the developmental trajectory of alveolar cells, we utilized STREAM (v1.1) [25] to approximate gene expression matrix in a structure called the principal graph, which is a set of curves that naturally describe the cell-pseudotime, trajectories, and branching points. In detail, we firstly constructed the principal graph through `seed_elastic_principal_graph(clustering="kmeans", n_clusters=12, use_vis=True)` based on recomputed original UMAP embedding, and then smoothed the trajectory through `elastic_principal_graph(epg_alpha=0.02, epg_mu=0.1, epg_lambda=0.01)`. Pseudotime heatmap was generated followed by the same protocol in airway trajectory.

To further verify the robustness of alveolar cells’s developmental trajectory, we applied scVelo (v0.2.5) [26] to recover the directed dynamic information by leveraging splicing kinetics, the result of velocity is highly consistent with the trajectory revealed by STREAM.

**Spatial cell-cell communication analysis**

For ST data, SpatialDM (v0.1.0) [22] was used to explore the spatially cell-cell communication across slices. We filtered out LRs of `CellChatDB.mouse` whose ligand and receptor are the same as each other and retained 1,998 valid LRs in three types of interactions – `Secreted Signaling`, `ECM-Receptor`, and `Cell-Cell Contact`. To calculate the proper distance matrix for different resolution, we modified parameters of `weight_matrix` based on maximum center-to-center ineraction distance (200 μm). `spatialdm_global(n_perm = 1000, method="z-score")` was used to identify significantly interacting LR pairs in whole slice, and then `spatialdm_local(n_perm = 1000, method="z-score")` was used to identify locally significant spots for each LR.

In total, SpatialDM provided us spatially signicificant LRs in each slice, which can be inspected manually based on image. To identitify cluster-specific LRs across slices, we applied differential analysis on each annotated cluster based on `-log10(local_z_p)`, and generated the cluster-spcific LRs's enrichment heatmap using SCP (v0.5.1). While the results of SpatialDM cannot accurately correspond to cell cluster derived from single-cell expression profile, we further performed cell-cell communication on the scRNA-seq reference to provide precise cell cluster information.

**scRNA cell-cell communication analysis**

For scRNA-seq data, CellChat (v1.6.1) [27] was used to explore the cell-cell communication within clusters. To maintain the consistency of cell-cell communication results between scRNA-seq and ST data, we applied the same LR database to CellChat. Firstly, we used `computeCommunProb(type = "triMean", trim = 0.1)` to calculate the communication probability between clusters, and then aggragate the results through `aggregateNet(thresh = 0.05)`.

**Single-cell-spatial cell-cell communication of alveolar microenvironment**

For cell clusters with discrete spatial distribution, we preferentially extract cluster-specific LRs based on Cellchat results. As for developmentally dynamic cell-cell communication in alveolar microenvironment, we firstly screened LRs specific to the alveolar microenvironment-related cell cluster using CellChat results, and then filtered out LRs which were not significant in CellChat and SpatialDM, and finally generated the COLLAGEN's enrichment heatmap using SCP (v0.5.1).

**Transposable Element (TE) quantification and analysis**

We utilized scTE (v1.0) [28] to quantifying TE expression from ST data. Firstly, we built the genome indice for scTE from gencode.M21, and then we put aligned sequence reads (.bam) into scTE for TE quantification. All steps are completed in accordance with the official tutorial (https://github.com/JiekaiLab/scTE). To identify cluster-specific TEs, we combined TE profile of different slices as the same protocol of ST data and then applied differential analysis on each annotated cluster based on TE expression. Finally we identified 240 cluster-specific TEs (126 of LTR, 66 of LINE, 28 of SINE, 17 of DNA, and 3 of Satellite) through `FindAllMarkers`.

**Analysis of alveolar niche before/after birth**

We selected alveolar regions of E18.5, P0 and P3 respectively according to the manual annotations above, and each region was about 10 x 10 spots. The principle was that most of the regions were annotated as cell types related to the alveolar microenvironment. In order to ensure the robustness of the analysis results, we selected 3 regions in each slice.

In order to identify the differential genes with gradient changes before and after birth in the alveolar regions, we performed differentially expressed genes (DEGs) analysis of E18.5 v.s. P0, E18.5 v.s. P3, and P0 v.s. P3. Then the intersection of each group of upregulated DEGs was defined as the upregulated gene set before birth; conversely, the intersection of each group of downregulated DEGs was defined as the upregulated gene set after birth.

**References**

[1] Carey MA, Card JW, Voltz JW, Germolec DR, Korach KS, Zeldin DC. The impact of sex and sex hormones on lung physiology and disease: lessons from animal studies. Am J Physiol Lung Cell Mol Physiol 2007;293:L272-8.

[2] Laube M, Thome UH. Y It Matters-Sex Differences in Fetal Lung Development. Biomolecules 2022;12.

[3] Liptzin DR, Landau LI, Taussig LM. Sex and the lung: Observations, hypotheses, and future directions. Pediatr Pulmonol 2015;50:1159-69.

[4] Syrett CM, Sierra I, Berry CL, Beiting D, Anguera MC. Sex-Specific Gene Expression Differences Are Evident in Human Embryonic Stem Cells and During In Vitro Differentiation of Human Placental Progenitor Cells. Stem Cells Dev 2018;27:1360-75.

[5] Liu Y, Yang M, Deng Y, Su G, Enninful A, Guo CC, et al. High-Spatial-Resolution Multi-Omics Sequencing via Deterministic Barcoding in Tissue. Cell 2020;183:1665-81.e18.

[6] Chen A, Liao S, Cheng M, Ma K, Wu L, Lai Y, et al. Spatiotemporal transcriptomic atlas of mouse organogenesis using DNA nanoball-patterned arrays. Cell 2022;185:1777-92.e21.

[7] Jiang F, Zhou X, Qian Y, Zhu M, Wang L, Li Z, et al. Simultaneous profiling of spatial gene expression and chromatin accessibility during mouse brain development. Nat Methods 2023;20:1048-57.

[8] Kechin A, Boyarskikh U, Kel A, Filipenko M. cutPrimers: A New Tool for Accurate Cutting of Primers from Reads of Targeted Next Generation Sequencing. J Comput Biol 2017;24:1138-43.

[9] Smith T, Heger A, Sudbery I. UMI-tools: modeling sequencing errors in Unique Molecular Identifiers to improve quantification accuracy. Genome Res 2017;27:491-9.

[10] Parekh S, Ziegenhain C, Vieth B, Enard W, Hellmann I. zUMIs - A fast and flexible pipeline to process RNA sequencing data with UMIs. Gigascience 2018;7.

[11] Choudhary S, Satija R. Comparison and evaluation of statistical error models for scRNA-seq. Genome Biol 2022;23:27.

[12] Hao Y, Hao S, Andersen-Nissen E, Mauck WM, 3rd, Zheng S, Butler A, et al. Integrated analysis of multimodal single-cell data. Cell 2021;184:3573-87.e29.

[13] Du Y, Kitzmiller JA, Sridharan A, Perl AK, Bridges JP, Misra RS, et al. Lung Gene Expression Analysis (LGEA): an integrative web portal for comprehensive gene expression data analysis in lung development. Thorax 2017;72:481-4.

[14] Sun X, Perl AK, Li R, Bell SM, Sajti E, Kalinichenko VV, et al. A census of the lung: CellCards from LungMAP. Dev Cell 2022;57:112-45.e2.

[15] Cable DM, Murray E, Zou LS, Goeva A, Macosko EZ, Chen F, et al. Robust decomposition of cell type mixtures in spatial transcriptomics. Nat Biotechnol 2022;40:517-26.

[16] Kleshchevnikov V, Shmatko A, Dann E, Aivazidis A, King HW, Li T, et al. Cell2location maps fine-grained cell types in spatial transcriptomics. Nat Biotechnol 2022;40:661-71.

[17] Negretti NM, Plosa EJ, Benjamin JT, Schuler BA, Habermann AC, Jetter CS, et al. A single-cell atlas of mouse lung development. Development 2021;148.

[18] Bolger AM, Lohse M, Usadel B. Trimmomatic: a flexible trimmer for Illumina sequence data. Bioinformatics 2014;30:2114-20.

[19] Yu W, Uzun Y, Zhu Q, Chen C, Tan K. scATAC-pro: a comprehensive workbench for single-cell chromatin accessibility sequencing data. Genome Biol 2020;21:94.

[20] Bravo González-Blas C, De Winter S, Hulselmans G, Hecker N, Matetovici I, Christiaens V, et al. SCENIC+: single-cell multiomic inference of enhancers and gene regulatory networks. Nat Methods 2023;20:1355-67.

[21] Van de Sande B, Flerin C, Davie K, De Waegeneer M, Hulselmans G, Aibar S, et al. A scalable SCENIC workflow for single-cell gene regulatory network analysis. Nat Protoc 2020;15:2247-76.

[22] Li Z, Wang T, Liu P, Huang Y. SpatialDM for rapid identification of spatially co-expressed ligand-receptor and revealing cell-cell communication patterns. Nat Commun 2023;14:3995.

[23] Qiu X, Mao Q, Tang Y, Wang L, Chawla R, Pliner HA, et al. Reversed graph embedding resolves complex single-cell trajectories. Nat Methods 2017;14:979-82.

[24] Gulati GS, Sikandar SS, Wesche DJ, Manjunath A, Bharadwaj A, Berger MJ, et al. Single-cell transcriptional diversity is a hallmark of developmental potential. Science 2020;367:405-11.

[25] Chen H, Albergante L, Hsu JY, Lareau CA, Lo Bosco G, Guan J, et al. Single-cell trajectories reconstruction, exploration and mapping of omics data with STREAM. Nat Commun 2019;10:1903.

[26] Bergen V, Lange M, Peidli S, Wolf FA, Theis FJ. Generalizing RNA velocity to transient cell states through dynamical modeling. Nat Biotechnol 2020;38:1408-14.

[27] Jin S, Guerrero-Juarez CF, Zhang L, Chang I, Ramos R, Kuan CH, et al. Inference and analysis of cell-cell communication using CellChat. Nat Commun 2021;12:1088.

[28] He J, Babarinde IA, Sun L, Xu S, Chen R, Shi J, et al. Identifying transposable element expression dynamics and heterogeneity during development at the single-cell level with a processing pipeline scTE. Nat Commun 2021;12:1456.
